# Maintenance of haematopoietic stem cells by JAK inhibition and increased tyrosine-unphosphorylated STAT5

**DOI:** 10.1101/2024.05.29.596460

**Authors:** Matthew J Williams, Xiaonan Wang, Hugo P Bastos, Gabriela Grondys-Kotarba, Carys Johnson, Nicole Mende, Emily F Calderbank, Michelle Wantoch, Hyun Jung Park, Qin Wu, Shucheng Jin, Giovanna Mantica, Rebecca Hannah, Nicola K Wilson, Dean C Pask, Tina L Hamilton, Sarah J Kinston, Ryan Asby, Rachel Sneade, Joanna Baxter, Peter Campbell, George S Vassiliou, Elisa Laurenti, Juan Li, Berthold Göttgens, Anthony R Green

## Abstract

Normal and malignant hematopoietic stem cells (HSCs) are controlled by extracellular cues including cytokine signalling through the JAK/STAT pathway. Here, we show that STAT5-deficient HSCs exhibit an unusual phenotype: while reduced multi-lineage repopulation and reduced self-renewal are commonly associated with overproliferation and exhaustion, they are instead associated with reduced cell-cycle progression and increased differentiation in STAT5-deficient HSCs. Mechanistic studies show that unphosphorylated-STAT5 (uSTAT5) contributes to this phenotype by constraining HSC differentiation, promoting HSC maintenance and upregulating transcriptional programs associated with stemness. The JAK1/2 inhibitor ruxolitinib increases levels of uSTAT5, constrains differentiation and proliferation of murine HSCs, promotes their maintenance and upregulates transcriptional programs associated with stemness. Ruxolitinib also enhances clonogenicity of normal human HSPCs, CALR-mutant murine HSCs and HSPCs from patients with myelofibrosis. Our results therefore reveal a previously unrecognized role for uSTAT5 in controlling HSC function, highlight JAK inhibition as a strategy for enhancing HSC function and provide insights into the failure of JAK inhibitors to eradicate myeloproliferative neoplasms.

## Introduction

Hematopoietic stem cells (HSCs) are a highly quiescent population of cells responsible for continued production of mature blood cells throughout life.^1,2^ Their ability to respond to environmental signals is important for maintaining homeostasis and for HSCs to respond to a variety of stresses.^3–6^

The JAK-STAT pathway regulates multiple developmental and adult stem cell populations^7–9^ and is dysregulated in a variety of haematological malignancies and other cancers.^10,11^ The signal transducer and activator of transcription 5 (STAT5) is an essential downstream mediator of cytokine signalling at multiple stages of haematopoiesis.^12–16^ In eutherian mammals, two closely related STAT5 isoforms,^17^ STAT5A and STAT5B, display distinct and redundant functions in different cell types.^18–21^ Mice lacking both genes, or the N-terminal domains of both genes, develop severe anaemia and leukopenia,^22–26^ associated with reduced survival and proliferation of erythroblasts.^15,16^ Conversely, high levels of STAT5 activity in haematopoietic stem and progenitor cells (HSPCs) drives erythroid differentiation.^27,28^

STAT5A and STAT5B contain critical regulatory tyrosine residues (Y694 and Y699 respectively) that are essential for activation of canonical pSTAT5 target genes.^29,30^ These residues are phosphorylated by Janus-kinases (JAKs)^31^ activated in response to multiple cytokines^3–6^ including IL-3^32^ and thrombopoietin (THPO).^33^ Tyrosine-phosphorylated STAT5 (pSTAT5) accumulates in the nucleus, binds to DNA and regulates transcription of target genes.^34^ STAT5 phosphorylation is transient as pSTAT5 rapidly promotes the expression of negative regulators of JAK-STAT signalling, including suppressors of cytokine signalling (SOCS), tyrosine phosphatases, and protein inhibitors of STATs.^35,36^

Elevated STAT5 phosphorylation is observed in many haematological malignancies^37,38^ and solid tumours.^39,40^ Activation of the JAK-STAT pathway is especially common in the myeloproliferative neoplasms (MPNs), >90% of which contain driver mutations which activate JAK-STAT signalling.^41–46^ JAK inhibitors are used to treat MPN patients with advanced disease^47^ but, although they can result in symptomatic improvement, they rarely reduce allele burden,^48–50^ suggesting that they fail to eradicate malignant HSCs.

Loss of both STAT5 genes results in reduced numbers of immunophenotypically-defined HSCs,^26,51,52^ as well as defective repopulation by foetal liver and adult bone marrow (BM).^26,53,54^ STAT5B is dominant in multipotent HPC7 cells^55^ and STAT5B-deficient, but not STAT5A-deficient, BM showed functional defects in serial transplants.^52^ However several aspects of STAT5 function in HSPCs remain unclear or have been the subject of conflicting reports: both increased^26,51,52^ and reduced cycling^56^ have been observed in HSPCs after STAT5 loss, while STAT5 phosphorylation is associated with increased proliferation.^57^ Moreover both STAT5 knockdown^55^ and constitutively active STAT5A overexpression^27,28^ have been reported to increase HSPC differentiation. Insight into at least some of these apparent paradoxes came from the demonstration that STAT5 lacking phosphorylation of its critical tyrosine (uSTAT5) is present in the nucleus of HSPCs and represses megakaryocytic differentiation by restricting access of megakaryocytic transcription factors to target genes.^55^ Cytokine-mediated phosphorylation of STAT5 therefore triggers two distinct transcriptional consequences: activation of a canonical pSTAT5-driven program that regulates proliferation and apoptosis; and loss of a uSTAT5 program that restrains megakaryocytic differentiation.

Given our limited understanding of the function of STAT5 in HSCs and the complete lack of information about the role of uSTAT5 in primitive HSCs, we addressed these issues using genetically modified mice combined with single cell approaches.

## Results

### STAT5 loss results in defective HSC function

Previous reports showed that STAT5^−/−^ foetal liver and adult BM cells displayed reduced repopulation in transplantation assays^26,54^ but it was unclear if this was a consequence of reduced HSC number or whether STAT5^−/−^ HSCs are also functionally impaired. We therefore crossed mice carrying a floxed *Stat5a/5b* allele^25^ with Mx1Cre mice and used Poly:IC to delete both *Stat5a* and *Stat5b* loci with ∼90% efficiency in haematopoietic cells (Fig. S1A-C).

Consistent with previous reports,^25,26^ STAT5 deletion resulted in anaemia, leukopenia and reduced BM cellularity (Fig. S1D, S1E). In STAT5-deficient BM the frequencies of immunophenotypic HSCs (both ESLAM; Lin^−^CD150^+^CD45^+^CD48^−^EPCR^+^, and LT-HSC; Lin^−^Sca1^+^cKit^+^CD150^+^CD48^−^CD34^−^Flk2^−^; Fig.Fig. 1A-B) and B-cells (Fig. 1C) were reduced and the proportion of erythroid progenitors (CFU-e; Lin^−^Sca1^−^cKit^+^CD41^−^ CD16/32^−^CD105^+^CD150^−^) was increased (Fig. 1D), but other mature and progenitor cell types were unaltered (Fig. S1F-I). In the spleen, STAT5 deletion reduced B-cell frequency (Fig. S1J) and increased frequencies of erythroid progenitors (CFU-e, PreCFU-e; Lin^−^Sca1^−^cKit^+^CD41^−^CD16/32^−^CD105^+^CD150^+^) and all stages of erythroblast differentiation (Fig. 1E-F).

**Fig. 1:**
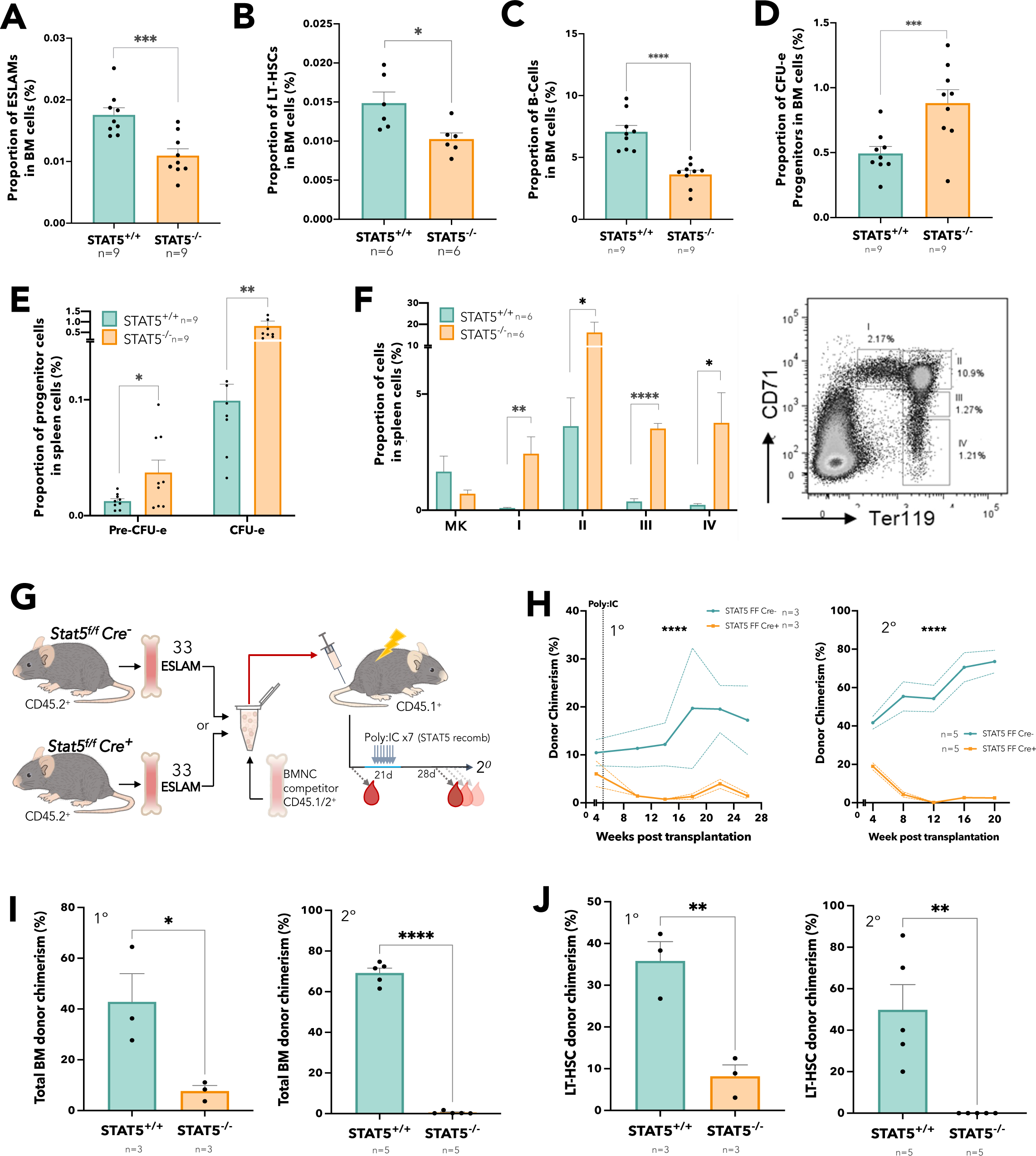
STAT5 loss results in defective HSC function. (A) Bar plot showing the frequency of ESLAM HSCs (CD45^+^CD150^+^CD48^−^EPCR^+^) in bone marrow mono-nuclear cells (BMMNCs) from WT and STAT5-deficientmice (mean ± SEM) (B) Bar plot showing the frequency of LT-HSCs (Lin^−^Sca1^−^c-Kit^+^CD150^+^CD48^−^CD34^−^Flk2^−^) in BMMNCs (mean ± SEM). (C) Bar plot showing the frequency of B-cells (B220^+^) in BMMNCs (mean ± SEM). (D) Bar plot showing the frequency of CFU-e progenitors (Lin^−^Sca1^−^cKit^+^CD41^−^CD16/32^−^CD105^+^CD150^−^) in BMMNCs (mean ± SEM). (E) Bar plots showing the frequency of CFU-e and pre-CFU-e (Lin^−^Sca1^−^cKit^+^CD41^−^CD16/32^−^CD105^+^CD150^+^) cells in spleen mono-nuclear cells (mean ± SEM). (F) Bar plots (left) showing the frequency of megakaryocyte (CD41^+^CD42^+^) and erythroid precursor cells (I, CD71^hi^Ter119^mid^; II, CD71^hi^Ter119^hi^; III, CD71^mid^Ter119^hi^; IV, CD71^low^Ter119^hi^) in spleen mono-nuclear cells (mean± SEM) with a representative flow cytometry plot (right) showing the gating of different stages of erythroid precursor cells in terminal differentiation. (G) Schematic diagram showing 33 FACS purified BM ESLAM HSCs were transplanted into irradiated recipient mice with 5×10^5^ competitor BMMNCs. STAT5 were deleted in Cre^+^ donor cells after transplantation by repeated injection (x7) with poly:IC in recipients. Blood was taken before and after STAT5 deletion and was followed for 5 months post deletion before serial transplantation of 3×10^6^ primary recipient BMMNCs. (H) Connected line graphs showing donor chimerism in peripheral blood mono-nuclear cells at each time point in primary and secondary recipients (mean ± SEM). Dotted line indicates initiation of Poly:IC injections. Asterisks indicate significant differences by ANOVA column factor (****, p<0.0001). (I) Bar plots showing total BMMNC donor chimerism in primary and secondary recipients (mean ± SEM). (J) Bar plots showing LT-HSC donor chimerism in primary and secondary recipients (mean ± SEM). Asterisks indicate significant differences by Student’s t test (****, p<0.0001; ***, p<0.001; **, p<0.01; *, p<0.05) unless otherwise indicated.

Droplet-based (10XGenomics) scRNAseq was performed to assess the haematopoietic stem/progenitor cell (HSPC) landscape. BM LK (Lin^−^cKit^+^) cells from pairs of STAT5^−/−^ and wildtype (WT) control mice were projected onto a previously published LK dataset^60^ and then a phenotypically-defined HSPC dataset,^61^ and cell types were annotated based on their nearest neighbours. Cells within the LT-HSC, ST-HSC, MPP, myeloid, early- and mid-erythroid clusters were relatively reduced in STAT5^−/−^ mice, while the abundance of cells within late-erythroid and lymphoid clusters were relatively increased (Fig. S1K). These results confirm and extend previous reports and show that STAT5 deficiency causes wide-spread alterations of haematopoietic progenitors including reduced numbers of HSCs.

In competitive transplantation experiments using highly purified ESLAM HSCs (Fig. 1G and Fig. S1L) STAT5-deficient HSCs displayed significantly reduced multilineage repopulation in blood (Fig. 1H, Fig. S1M-O) and BM (Fig. 1I, Fig. S1P) of primary recipients. There was almost no repopulation of blood or BM in secondary recipients. Few or no STAT5-deficient LT-HSCs were observed in the BM of primary or secondary recipients respectively (Fig. 1J and Fig. S1Q). These data demonstrate that STAT5-deficient HSCs are not merely reduced in number but are also functionally impaired and display markedly reduced multi-lineage repopulation and self-renewal.

### STAT5-deficient HSCs display reduced cell cycle progression, increased differentiation, and reduced generation of lineage-negative progeny

To explore the molecular basis for HSC dysfunction, plate-based scRNAseq was performed on WT and STAT5-deficient ESLAM HSCs (Fig. S2A-C) and identified 308 differentially expressed genes (adj. p<0.01, LogFC>±0.5, Fig. S2D and Table S1), including canonical STAT5 targets (eg *Cish*, *Socs2* and *Bcl6*; Fig. S2E).

Gene set enrichment analysis identified 12 signatures enriched in STAT5-deficient HSCs (FDR<0.25; Table S2), including Wnt, Hedgehog and Kras pathways, together with 35 signatures that were depleted (FDR<0.25; Table S2), including JAK-STAT signalling, DNA repair and unfolded protein response. The most significantly depleted gene sets were cell cycle related signatures including E2F targets and DNA replication (Fig. 2A and Fig. S2F). Consistent with this observation, analysis of our separate 10X LK cell datasets showed that, compared to WT controls, far fewer STAT5-deficient LT-HSCs were in cycle (8.58 vs 2.82%, Fig. 2B). A less pronounced reduction in cell cycling was seen in STAT5-deficient ST-HSCs and MPPs.

**Fig. 2:**
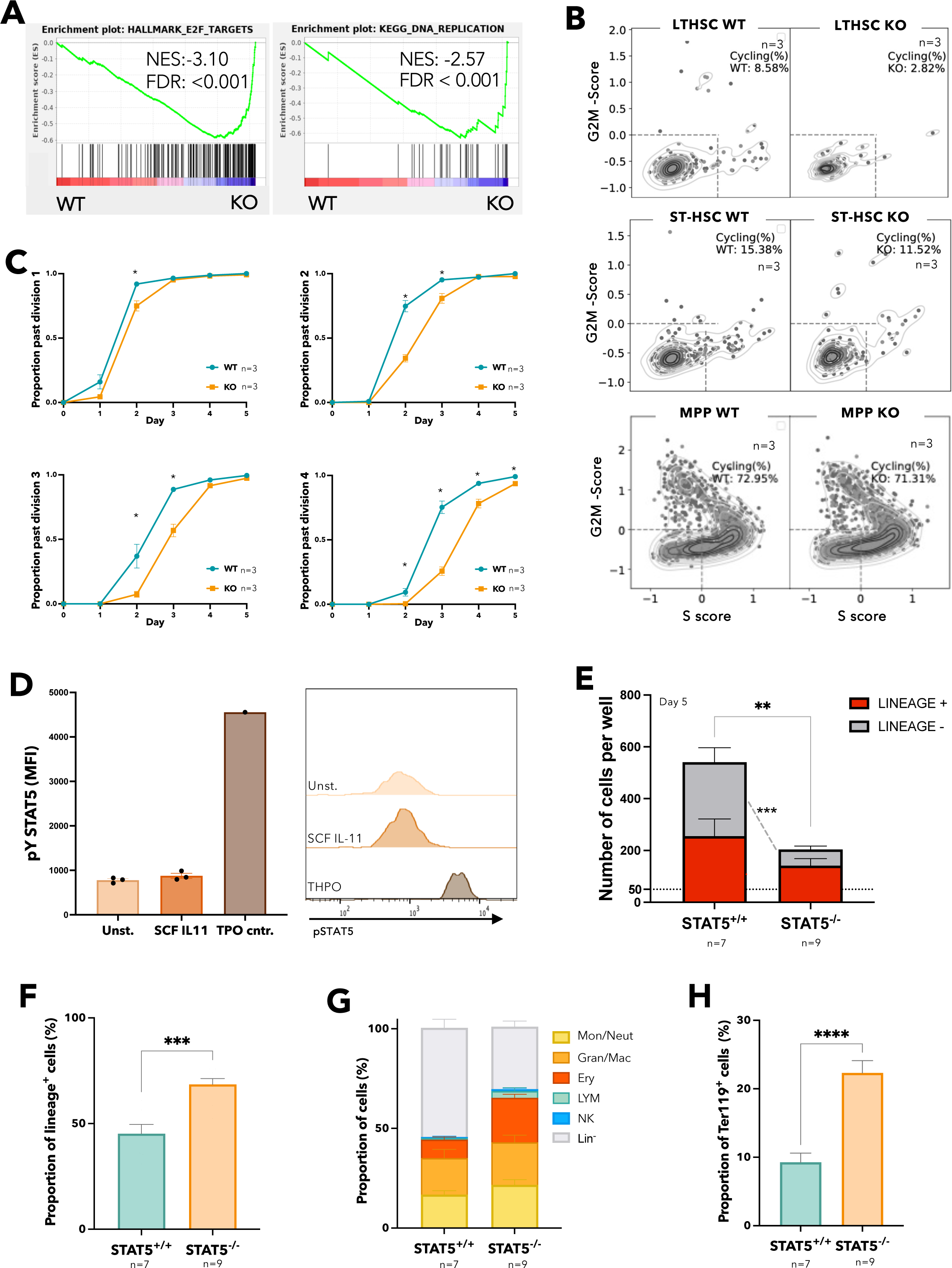
STAT5-deficient HSCs display reduced cell cycle entry, increased differentiation, and reduced retention of lineage-negative progeny. (A) Gene set enrichment analysis (GSEA) plots showing depleted cell cycle related signatures in STAT5-deficient ESLAM HSCs. ScRNAseq analysis using Smartseq platform was performed on FACS isolated ESLAM HSCs from STAT5^f/f^ Cre- or STAT5^f/f^ Cre+ bone marrow, 132 STAT5-deficient and 132 WT HSCs passed quality control and were used for downstream analysis. Normalised enrichment scores (NES) and false discovery rate (FDR) are indicated. (B) Plots showing cell cycle scores of LT-HSCs, ST-HSCs and MPPs that were transcriptionally isolated from scRNAseq datasets of WT and STAT5-deficient BM LK cells (STAT5 WT, n=3; STAT5KO, n=3). (C) Line graphs showing proportion of ESLAM HSCs that past first, second, third and fourth divisions at given timepoints (y axis) in single cell *in vitro* analysis (mean ± SEM). Results from 3 biological replicates across 3 experiments. (D) Bar plots (left) showing the mean fluorescent intensity (MFI) of pSTAT5 antibody staining of ESLAM HSCs by intracellular flow-cytometry analysis in unstimulated, maintenance culture conditions^62^ (SCF/IL-11), or TPO (200ng/mL) positive control conditions (mean ± SEM). Right; representative histogram of the intracellular flow-cytometry analysis showing the intensity of pSTAT5 staining in each condition. Results from three biological replicates. (E) Bar plot showing the number of cells in each well at day 5, from an initial culture of 50 ESLAM HSCs in SCF/IL11 maintenance condition. The number of cells expressing mature lineage markers (Ter119^+^, Ly6g^+^, CD11b^+^, NK1.1^+^, B220^+^, CD19^+^ or CD3e^+^) are shaded in red; the number of lineage negative cells are shaded in grey (mean ± SEM). Results from 9-7 biological replicates across 4 experiments. (F) Bar plot showing the proportion of cells expressing mature lineage markers after 5 days in culture originating from 50 ESLAM HSCs (mean ± SEM). (G) Bar plot showing the proportion of cells expressing specific mature lineage markers for monocytes and granulocytes (Ly6g^+^), granulocytes and macrophages (CD11b^+^), erythroid (Ter119^+^), lymphocytes (CD3e^+^/CD19^+^/B220^+^) and natural killer cells (NK1.1^+^) after 5 days in culture originating from 50 ESLAM HSCs (mean ± SEM). (H) Bar plot showing the frequency of Ter119^+^ cells after 5 days in culture originating from 50 ESLAM HSCs (mean ± SEM). Asterisks indicate significant differences by Student’s t test (****, p<0.0001; ***, p<0.001; **, p<0.01; *, p<0.05).

Ki-67/DAPI staining and flow cytometry indicated that the fraction of cells in G_0_, G_1_ and S-G_2_-M were comparable between WT and STAT5-deficient ESLAM HSCs (Fig. S2G-2I). Ki-67/DAPI analysis represents a snapshot, which does not capture subtle but relevant changes in quiescence maintenance. We therefore measured the division kinetics of single HSCs (as previously described^62^). STAT5-deficient HSCs were indeed slower to enter their first and subsequent divisions (Fig. 2C), thus demonstrating that STAT5 is required for normal HSC cell cycle progression.

The functional consequences of STAT5 deficiency described above reflect the combined effect of losing both pSTAT5 and uSTAT5. To identify those consequences attributable to loss of uSTAT5, we grew HSCs in SCF and IL-11 media (previously described to maintain HSCs^62^), in which pSTAT5 was undetectable in HSCs (Fig. 2D). After 5 days in this culture condition, STAT5-deficient HSCs produced fewer cells overall, with markedly fewer lineage-negative cells (Fig. 2E) and an increase in the proportion of lineage-positive cells (Fig. 2F). The proportion of each lineage tested increased (Fig. 2G, Fig. S2J-M) with the erythroid lineage (Ter119^+^) reaching statistical significance (Fig. 2H). Similar results were obtained with HSCs cultured for 4 or 6 days (Fig. S2N) with no difference in the frequency of apoptotic cells (Fig. S2O). These data indicate that loss of uSTAT5 is responsible for increased HSC differentiation and reduced generation of lineage-negative cells.

Together our results therefore demonstrate that STAT5 loss results in an unusual HSC phenotype consisting of reduced cell cycle progression and yet increased differentiation.

### Unphosphorylated STAT5 constrains HSC differentiation and upregulates transcriptional programs associated with HSC maintenance

To further explore the role of uSTAT5 in HSCs we utilised lentiviral expression approach. STAT5B is the dominant form of STAT5 protein in multipotent HPC7 cells^55^ and long-term repopulating HSCs.^52^ STAT5B-Y699F (STAT5-YF), which prevents phosphorylation at this critical residue, was introduced into STAT5^+/+^ or STAT5^−/−^ HSCs along with empty vector (EV) controls (Fig. 3A). STAT5-YF and EV constructs showed comparable expansion and survival in STAT5^+/+^ HSCs and also in STAT5^−/−^ HSCs (Fig. S3A-B). When introduced into either STAT5^+/+^ or STAT5^−/−^ HSCs, STAT5-YF resulted in reduced differentiation (Fig. 3B). These observations accord with our studies of STAT5^−/−^ HSCs, which indicated that loss of uSTAT5 enhances their differentiation (see above). Thus, both gain-of-function and loss-of-function approaches indicate that uSTAT5 constrains HSC differentiation. STAT5-YF expression increased total STAT5 levels 2-3 fold (Fig. S3C-D) and so our results indicate that the functional consequences of STAT5-YF reflect a ∼2-3 fold increase in uSTAT5, given that the vast majority of potentially phosphorylatable STAT5 (ie can be phosphorylated by THPO) remains unphosphorylated in HSCs cultured in SCF/IL-11 (Fig. 2D and Fig. S3E).

**Fig. 3:**
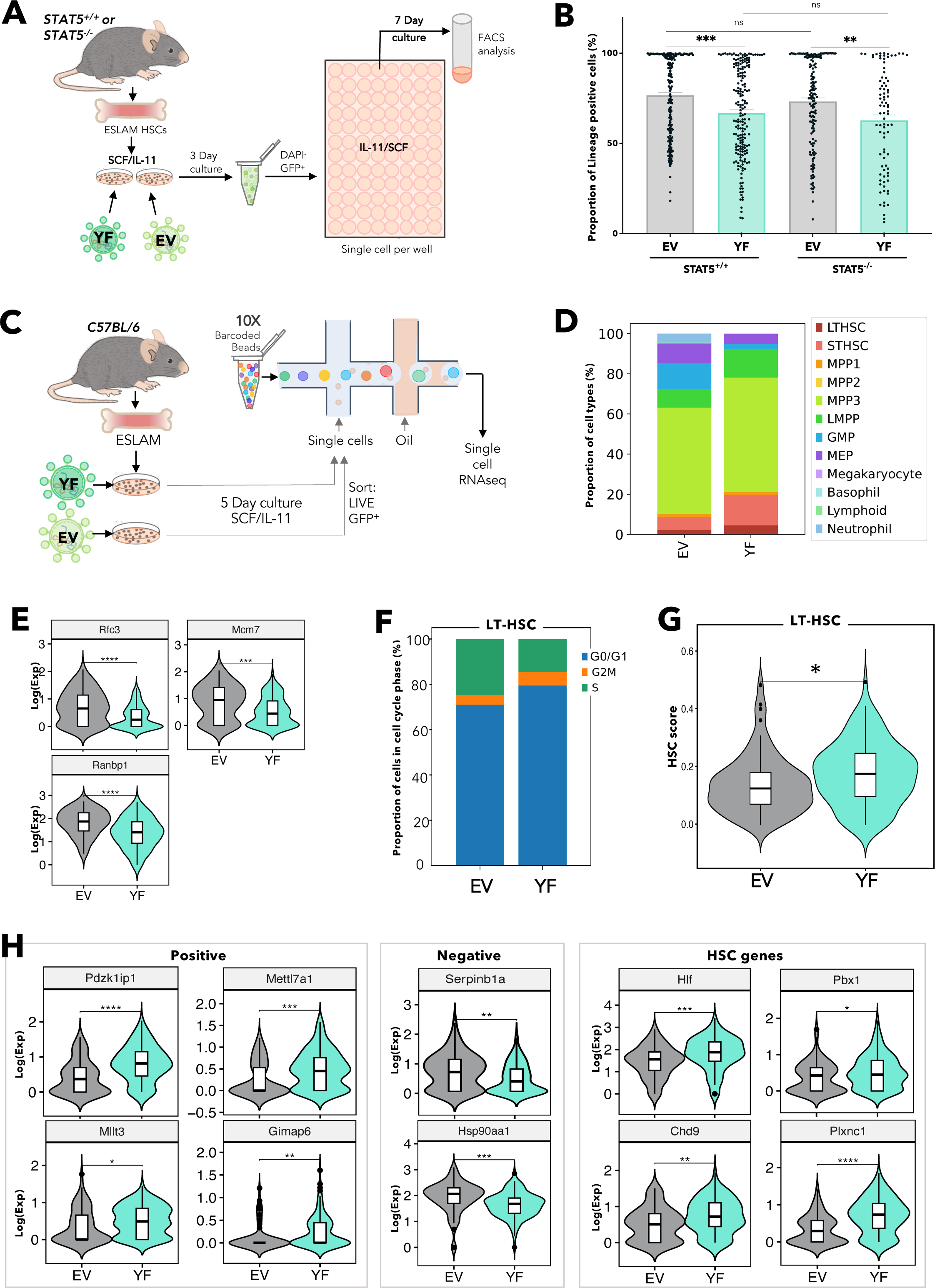
Unphosphorylated STAT5 constrains HSC differentiation and upregulates transcriptional programs associated with HSC maintenance. (A) Schematic diagram showing the experimental outline of the ex vivo functional analysis of STAT5-deficient or WT ESLAM HSCs that were transduced with lentivirus containing STAT5B-Y699F (YF) or EV in maintenance cultures.^62^ After three days of transduction, GFP^+^ living cells were FACS sorted into single cell assays. (B) Bar plots showing the proportion of cells expressing mature lineage markers (Ter119^+^/Ly6g^+^/CD11b^+^/B220^+^/CD3e^+^). Each dot represents a single clone and bars represent mean lineage positive marker frequency (± SEM). Asterisks indicate significant differences by Student’s t test (***, p<0.001; **, p<0.01). Results from 6 to 5 independent biological replicates across 5 experiments in STAT5^+/+^ settings and 4 to 3 independent biological replicates across 3 experiments in STAT5^−/−^ settings. (C) Schematic diagram showing the outline of scRNAseq of WT ESLAM HSCs that were transduced with lentivirus containing STAT5B-Y699F or EV in maintenance cultures^62^ and were allowed to expand for 5 days. GFP^+^ living cells were then sorted for 10X Genomics scRNAseq. (D) Bar plots showing the proportion of annotated cell types in GFP^+^ HSC derived cultures after 5-days in SCF/IL-11 cultures; single cells were projected onto a previously published scRNAseq dataset of LK HSPC cells^60^ and then a phenotypically-defined HSPC dataset,^61^ and cell types were annotated based on their nearest neighbours to ascribe cell identity and cell type annotation. Results from 2 independent biological replicates in 2 experiments. (E) Violin plots showing significantly differentially expressed genes associated with cell cycle regulation, in transcriptionally isolated LT-HSCs (STAT5B-YF, n=83; EV, n=53). (F) Bar plots showing the frequency of LT-HSCs in each phase (G_0_/G_1_ phases, reflecting cells that scored below 0 for S-phase or G_2_M cell cycle scores; S or G_2_M phases) of the cell cycle based on transcriptional cell cycle scores. (G) Violin plot showing the geometric mean distribution of HSC scores in LT-HSCs expressing STAT5B-YF or EV. The HSC score was calculated using the HSC score tool that identifies potential mouse bone marrow HSCs from scRNA-Seq data.^63^ This tool considers the expression of genes that are either positively or negatively corelated with HSC long-term repopulating capacity.^64^ (H) Violin plots showing significantly differentially expressed genes that are positively associated with functional long-term repopulating HSCs (*Pdzk1ip1*, *Mettl7a1*, *Mllt3*, *Gimap1*), negatively associated with functional long-term repopulating HSCs (*Serpinb1a*, *Hsp90aa1*), or genes with reported functions in maintaining HSCs (*Hlf*, *Chd9*, *Pbx1*, *Plxnc1*). All data is combined from two independent experiments. Asterisks indicate significant differences by Student t test (****, p<0.0001; ***, p<0.001; **, p <0.01; *, p <0.05).

The transcriptional consequences of STAT5-YF expression in ESLAM HSCs were explored using 10X Genomics scRNAseq (Fig. 3C). Since STAT5^−/−^ and STAT5^+/+^ HSCs responded similarly to STAT5-YF overexpression and STAT5-deficient HSCs are less abundant, STAT5^+/+^ HSCs were used for this analysis. *Stat5b* transcripts increased 2-fold in STAT5-YF infected cells (Fig. S3F) consistent with protein levels (Fig. S3B). Infected cells were projected onto a previously published scRNAseq dataset of LK cells^60^ and then a phenotypically-defined HSPC dataset,^61^ and cell types were annotated based on their nearest neighbours. Compared to control EV cultures, STAT5-YF cultures contained fewer differentiated cell types (eg GMPs, MEPs, and neutrophils) but more early stem/progenitor cells (LT-HSCs and ST-HSCs; Fig. 3D and Fig. S3G). These results accord well with our functional evidence that STAT5-YF constrains differentiation.

Within transcriptionally defined LT-HSCs, STAT5-YF expression was associated with upregulation/downregulation of 321/120 genes respectively (. S3H, Table S3). Canonical pSTAT5 target genes were unaffected by STAT5-YF expression (Fig. S3I), suggesting uSTAT5 expression does not repress pSTAT5 activity in a dominant-negative manner. Multiple cell-cycle genes (eg *Mcm7, Rfc3, Ranbp1*) were down-regulated in STAT5-YF infected HSCs (Fig. 3E) and fewer STAT5-YF HSCs were in S phase (Fig. 3F), collectively indicating that STAT5-YF expression was associated with increased HSC quiescence.

Moreover, compared to EV expressing HSCs, STAT5-YF expressing HSCs exhibited higher HSC scores (Fig. 3G) using a previously described algorithm that identifies durable long-term repopulating HSCs^63^ and takes into account the expression of genes that correlate either positively or negatively with HSC function.^64^ STAT5-YF HSCs also exhibited higher HSC scores using two other published HSC signatures (Fig. S3J).^65,66^ Indeed, positively-associated HSC score genes were upregulated in STAT5-YF HSCs, while anti-correlated genes were downregulated, and other genes reported to promote HSC maintenance were also upregulated (Fig. 3H). Together our transcriptional data therefore demonstrate that STAT5-YF regulates transcriptional networks associated with restrained differentiation, reduced cell cycling and increased HSC maintenance.

### Unphosphorylated STAT5 enhances HSPC clonogenicity *in vitro* and HSC maintenance *in vivo*

We next explored the effect of STAT5-YF expression on HSC function (Fig. 4A). In serial colony replating assays STAT5^+/+^ HSCs expressing STAT5-YF displayed enhanced colony generation in 4 independent experiments (Fig. 4B and Fig. S4A) demonstrating that uSTAT5 is sufficient to enhance generation of clonogenic progeny by WT HSCs. Introduction of STAT5-YF had no effect on the replating of STAT5^−/−^ HSCs, but these cells produced far fewer colonies for a shorter duration than WT cells (Fig. 4C and Fig. S4B) indicating a requirement for pSTAT5 in the replating assay, likely through its role driving programs associated with proliferation.^55^ Indeed, STAT5 phosphorylation was readily detectable in HSCs cultured in the replating assay media which contained IL-3 and IL-6 (Fig. S4C).

**Fig. 4:**
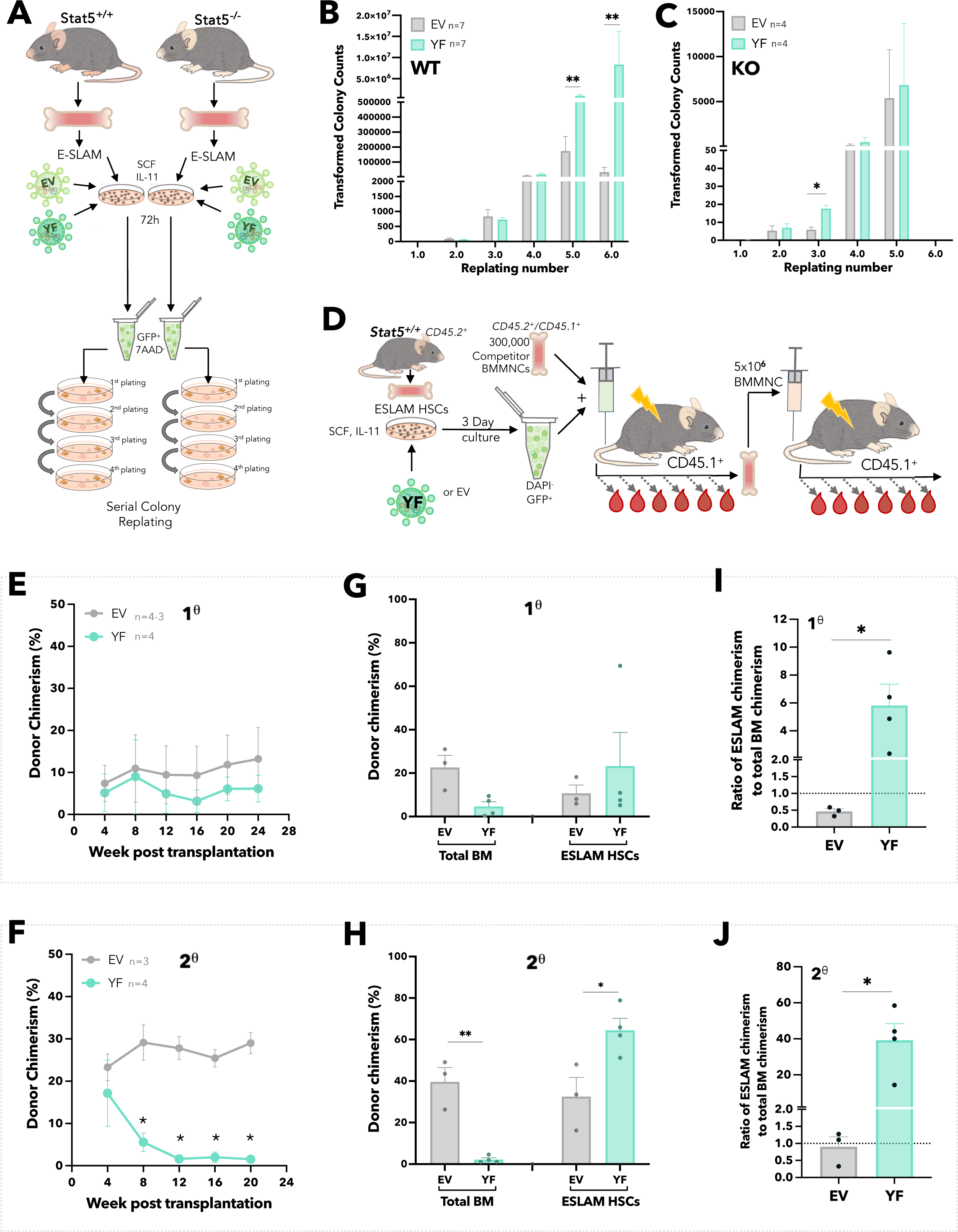
Unphosphorylated STAT5B enhances HSPC clonogenicity *in vitro* and HSC maintenance *in vivo*. (A) Schematic diagram showing the experimental outline of serial colony replating assays of STAT5-deficient or WT ESLAM HSCs that were transduced with lentivirus containing STAT5B-Y699F (YF) or EV in SCF/IL-11 maintenance cultures. After three days of transduction, GFP^+^ living cells were sorted for serial colony replating assays. (B) Bar plots showing the transformed colony numbers derived from WT HSPCs transduced with YF or EV lentivirus. Transformed colony counts = ((colony number x dilution factor) ÷ starting number of HSCs) (mean ± SEM). Results were from four independent experiments and 7 biological replicates. Asterisks indicate significant differences by Mann-Whitney t test (**, p<0.01). (C) Bar plots showing the transformed colony numbers of STAT5-deficient HSPCs transduced with YF or EV lentivirus. Transformed colony counts = ((colony number x dilution factor) ÷ starting number of HSCs)) (mean ± SEM). Results were from three independent experiments and 4 biological replicates. Asterisks indicate significant differences by Mann-Whitney t test (*, p<0.05). (D) Schematic diagram showing outline of the *in vivo* functional analysis of WT ESLAM HSCs that were transduced with lentivirus containing STAT5B-Y699F (YF) or EV. FACS sorted WT ESLAM HSCs (CD45.2^+^) were infected with lentivirus and cultured for 3 days in maintenance cultures, then equal number of GFP^+^ cells were FACS sorted (112 GFP^+^ cells/recipient) and injected into irradiated recipients (CD45.1^+^) with 3×10^5^ competitor BMMNCs (CD45.1^+^/CD45.2^+^). Donor chimerism was monitored every 28 days for over 6 months. Secondary transplantation was then performed using 5×10^6^ BMMNCs from primary recipients. (E) Connected line graph showing donor chimerism in primary recipients (mean ± SEM); experiment described in 4D. Chimerism was derived as the ratios of donor/(donor+competitor). (F) Connected line graph showing donor chimerism in secondary recipients (mean ± SEM); experiment described in 4D. Chimerism was derived as the ratios of donor/(donor+competitor). (G) Bar plots showing donor chimerism in total BMMNCs and ESLAM HSCs in BM of the primary transplant recipients (mean ± SEM). (H) Bar plots showing donor chimerism in total BMMNCs and ESLAM HSCs in BM of the secondary transplant recipients (mean ± SEM). (I) Bar plots showing the ratio of ESLAM HSC donor chimerism to total BMMNC chimerism in primary recipient BM (mean ± SEM; dotted line indicating 1:1 ratio). (J) Bar plots showing the ratio of ESLAM HSC donor chimerism to total BMMNC chimerism in secondary recipient BM (mean ± SEM; dotted line indicating 1:1 ratio). Chimerism was derived as the ratios of donor/(donor+competitor). Asterisks indicate significant differences by Student’s t test (**, p<0.01; *, p<0.05).

In competitive transplantation experiments (Fig. 4D), compared to control HSCs infected with EV, those carrying STAT5-YF generated peripheral blood donor chimerism that was modestly reduced in primary recipients (Fig. 4E) and dramatically reduced in secondary recipients (Fig. 4F, Fig. S4D). Furthermore, compared to control EV-infected HSCs, HSCs infected with STAT5-YF gave rise to reduced total BM chimerism but increased HSC chimerism in primary recipients (Fig. 4G) an observation that was even more striking in secondary recipients (Fig. 4H). Within individual primary recipients, the ratio of HSC chimerism to that individual’s total BM chimerism was substantially higher for mice that had received STAT5-YF HSCs compared to those that had received EV HSCs (Fig. 4I). This pattern was even more striking in secondary transplant recipients (Fig. 4J).

Together our results therefore indicate that STAT5-YF expression enhances the generation of clonogenic progeny by HSCs *in vitro* and reduces exit from the stem cell compartment i*n vivo*.

### Ruxolitinib enhances HSPC clonogenicity and maintains transplantable HSCs

JAK inhibitors, such as ruxolitinib, are predicted to increase the ratio of uSTAT5 to pSTAT5. Indeed, ruxolitinib treatment of cells with activated JAK/STAT signalling (driven by mutant JAK2 or mutant CALR), resulted in a dramatic reduction in pSTAT5 without a fall in total STAT5 protein levels (Fig. S5A). In primary HSCs levels of pSTAT5 (but not pSTAT1 or pSTAT3), were induced 4-fold in a ruxolitinib-sensitive manner when exposed to SCF, IL3 and IL6 (Fig. 5A, 5B and Fig. S5B-C). Total STAT5 protein levels were unchanged (Fig. 5C) and pSTAT5 target genes such as *Cish* and *Pim1* were downregulated in HSCs exposed to ruxolitinib (Fig. S5D).

**Fig. 5:**
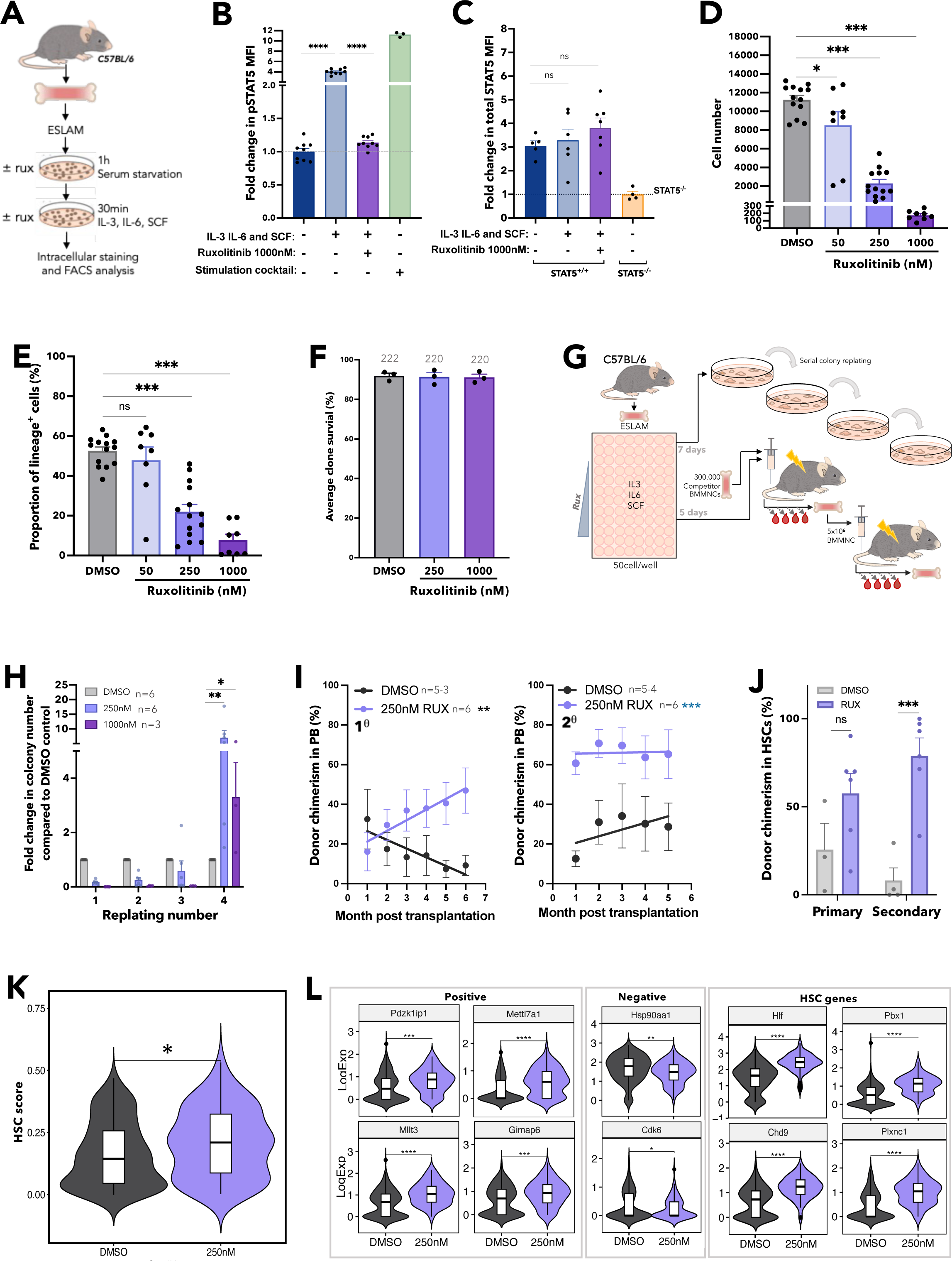
Ruxolitinib enhances HSPC clonogenicity and maintains transplantable HSCs. (A) Schematic diagram showing intracellular flow cytometric analysis of STAT5 proteins in ruxolitinib (RUX) treated WT ESLAM HSCs. WT ESLAM HSCs were sorted into serum starved media and starved for 1h before a 30-minute stimulation with complete medium containing IL-3, IL-6 and SCF in the presence or absence of RUX^89^, a stimulation cocktail containing THPO, Flt3-L and IFNα for positive control was included. Cells were then fixed and stained for intracellular flow cytometry. (B) Bar plots showing the mean fluorescent intensity (MFI) of pSTAT5 antibody staining in ESLAM HSCs described in ‘A’ normalised to the unstimulated condition which is indicated with dotted line (mean ± SEM). Each dot represents the mean fluorescent intensity of ESLAMs from a single mouse. Results from 3 independent experiments. (C) Bar plots showing the mean fluorescent intensity of total-STAT5 (tSTAT5) antibody staining in ESLAM HSCs, described in ‘A’ normalised to STAT5-deficient HSPCs which is indicated with dotted line (mean ± SEM). Results from 3 independent experiments. (D) Bar plots showing cell number per well in HSC derived cultures at each dose of ruxolitinib or vehicle after 7 days (mean ± SEM). 50 ESLAMs were seeded per well in 96-well plates in IL-3/IL-6/SCF^89^ cultures and were treated with DMSO or indicated doses of ruxolitinib. Results from 6 independent experiments. (E) Bar plot showing the proportion of cells expressing lineage positive markers (Ter119^+^/Ly6g^+^/CD11b^+^/B220^+^/CD3e^+^) after 7 days in culture at different concentrations (nM) of ruxolitinib (mean ± SEM). 50 ESLAMs were seeded per well in 96-well plates in IL-3/IL-6/SCF^89^ cultures and were treated with indicated doses of ruxolitinib. Results from 6 independent experiments. (F) Bar plots showing the clone survival rate of single HSCs after 5 days in culture. Single ESLAM HSCs were sorted per well and treated with vehicle or ruxolitinib, Clone survival rate was the proportion of wells that contained cells at day 5. Each dot represents the frequency of surviving clones from each of three independent experiments; bars show the mean ± SEM. (G) Schematic diagram showing serial colony replating assays and *in vivo* functional analysis for ESLAM HSCs treated with RUX or vehicle. 50 WT ESLAM HSCs were sorted per well into complete media^89^ with scaled doses of ruxolitinib or vehicle. Cells were harvested after 7 days and plated into serial colony replating assays. ESLAM HSC (CD45.2^+^) derived cells after 5 days in culture were harvested and transplanted into lethally irradiated recipient mice (CD45.1^+^) with 3×10^5^ fresh BMMNCs from competitor mice (CD45.1^+^/CD45.2^+^). Blood was analysed every 28 days for 6 months. Secondary transplants were then set up by transplanting 3×10^6^ bone marrow cells from the primary transplants recipients. (H) Bar plots showing the number of colonies produced by HSC-derived cultures treated with vehicle or ruxolitinib (250nM or 1000nM) for 7 days, normalised to the number of colonies produced by vehicle treated cultures at each week of replating. Results are shown as mean ± SEM and were from five independent experiments, three of which included 1000nM. Asterisks indicate significant differences by Mann-Whitney t test (**, p<0.01; *, p<0.05). (I) Scatter dot plot with linear regression line of best fit showing the peripheral blood donor chimerism in primary (left) and secondary (right) recipients transplanted with 5-day *ex vivo* cultured HSCs with ruxolitinib or vehicle. 50 ESLAMs from WT mice were seeded per well in IL-3/IL-6/SCF culture conditions and given DMSO or 250nM of ruxolitinib for 5 days before the cells were harvested and pooled for each condition and an equivalent of 10 starting ESLAMs was transplanted per recipient with 3×10^5^ competitor bone marrow cells. Each dot indicates mean donor chimerism and are shown as mean ± SEM. Black asterisks indicate significant differences in the slopes of linear regression modelling comparing chimerism of ruxolitinib treated donor cell to DMSO treated donor cell chimerism in the primary recipients (**, p<0.01). Blue asterisks indicate significant differences in y-intercepts of linear regressions modelling comparing chimerisms of ruxolitinib treated donor cell compared to DMSO treated donor cell in secondary transplants (***, p<0.001). (J) Bar plots showing the donor chimerism within the ESLAM HSC compartment at the end of primary and secondary recipients of 5-day *ex vivo* cultured HSCs with ruxolitinib or vehicle. Data are shown as mean ± SEM. (K) Violin plot showing the geometric mean distribution of HSC scores in LT-HSCs from 10x scRNAseq dataset of the cells treated with ruxolitinib or DMSO. The scores were calculated using the HSCscore tool that identifies potential mouse bone marrow HSCs from scRNA-Seq data.^63^ This tool considers the expression of genes that are either positively or negatively correlated with HSC long-term repopulating capacity.^64^ (L) Violin plots showing significantly differentially expressed genes that are positively associated with functional long-term repopulating HSCs (*Pdzk1ip1*, *Mettl7a1*, *Mllt3*, *Gimap6*), negatively associated with functional long-term repopulating HSCs (*Hsp90aa1*, *Cdk6*), or genes with reported functions in maintaining HSCs (*Hlf*, *Pbx1*, *Chd9*, *Plxnc1*). All data is combined from two independent experiments. Asterisks indicate significant differences by Student’s t test (****, p<0.0001; ***, p<0.001; **, p<0.01; *, p<0.05) unless otherwise indicated.

Ruxolitinib reduced, in a dose-dependent manner, the progeny generated by HSCs (Fig. 5D, Fig. S5E-F) and the proportion of lineage-positive cells (Fig. 5E, Fig. S5G-H). Two other JAK inhibitors, fedratinib and momelotinib, similarly reduced the expansion and differentiation of HSCs in culture (Fig. S5I-J). Treatment with ruxolitinib was not accompanied by reduced HSPC viability; even single ESLAM HSCs cultured with high doses of ruxolitinib (eg 1000nM, well above the therapeutic range) showed no difference in the proportion of wells containing one or more viable cells at 5 days (Fig. 5F). Moreover, treatment of lineage-depleted BM cells with ruxolitinib over-night resulted in apoptosis of mature cell types but had little effect on Lin^−^c-Kit^+^ cells suggesting that ruxolitinib does not affect survival of early HSPCs (Fig. S5K).

To investigate the effect of ruxolitinib on HSC function, serial colony replating assays and competitive transplants were performed (Fig. 5G). Compared to vehicle-treated HSCs, those exposed to ruxolitinib formed significantly more colonies in the final week of replating assays (Fig. 5H, Fig. S5L-M) indicating that ruxolitinib increased maintenance of clonogenic HSPCs in precultures. In two independent competitive repopulation experiments, vehicle-treated control cells gave rise to donor peripheral blood chimerism that gradually fell over the 5-month study period (Fig. 5I, Fig. S5N) as previously reported for cultured HSC donors.^67^ In marked contrast ruxolitinib-treated HSCs gave rise to levels of donor peripheral blood chimerism that were initially lower than controls and then were maintained or increased. In secondary recipients, donor HSCs originally treated with ruxolitinib displayed significantly higher peripheral blood chimerism (Fig. 5I, Fig. S5O). Moreover, primary recipient mice that had received HSCs precultured with ruxolitinib displayed increased chimerism in the HSC compartment, an effect that was even more marked in secondary recipients (Fig. 5J). Single cell RNAseq was used to explore the transcriptional consequences of ruxolitinib (Fig. S5P). Ruxolitinib-treated HSCs exhibited reduced expression of canonical pSTAT5 target genes (Fig. S5Q) and contained more transcriptionally defined LT-HSCs and ST-HSCs (Fig. S5R). Ruxolitinib-treated HSCs also showed increased HSC scores (Fig. 5K, Fig. S5S) using the 3 different methods^63,65,66^ that had also demonstrated increased HSC scores for STAT5-YF treated HSCs (Fig. 3G, Fig. S3J). Ruxolitinib-treated HSCs also showed increased scores for a signature derived by comparing STAT5-YF expressing HSCs to EV-transduced controls (Fig. S5T). Furthermore, ruxolitinib increased the expression of positively-associated HSC-score genes, reduced the expression of negatively-associated HSC-score genes and increased the expression of multiple other genes associated with HSC maintenance (Fig. 5L) in a manner similar to STAT5-YF expression (Fig. 3H). Several of these genes (eg *Pdzk1ip1, Gimap6, Hlf, Plxnc1* and *Chd9*) had previously been identified by ChIP studies^55^ as direct targets of uSTAT5 (Fig. S5U).

Together our data demonstrate that ruxolitinib pre-treatment enhanced HSPC clonogenicity and also the maintenance of transplantable HSCs. Moreover, the transcriptional consequences of ruxolitinib closely paralleled those observed for STAT5-YF expressing HSCs (Fig. 3G-H) indicating that the effects of ruxolitinib are mediated, at least in part, by uSTAT5.

### Ruxolitinib maintains murine and human myeloproliferative neoplasm HSPCs

Ruxolitinib alleviates symptoms, reduces splenomegaly and modestly extends overall survival in a subset of MPN patients with advanced disease.^47–49^ However it has little or no effect on allele burden and disease progression^48,49^ suggesting that ruxolitinib does not eradicate malignant HSCs. This has been attributed to ruxolitinib having a narrow therapeutic window as a consequence of dose limiting toxicity.^68,69^ However our data raise the possibility that JAK inhibitors might also inherently promote maintenance of mutant HSCs by increasing levels of uSTAT5.

We therefore studied the effect of ruxolitinib on CALR-mutant HSCs derived from a knock-in mouse model^58^ carrying a CALR-52bp deletion mutation commonly observed in human MPN patients^43^ and known to activate JAK/STAT signalling^70^ (Fig. 6A). Ruxolitinib reduced the number of progeny generated by CALR-mutant HSCs and also the proportion of lineage-positive cells (Fig. 6B, 6C). Ruxolitinib pre-treatment also enhanced the replating capacity of HSPCs derived from CALR-mutant HSCs (Fig. 6D, 6E, Fig. S6A-B) demonstrating that ruxolitinib maintains clonogenic HSPCs.

**Fig. 6:**
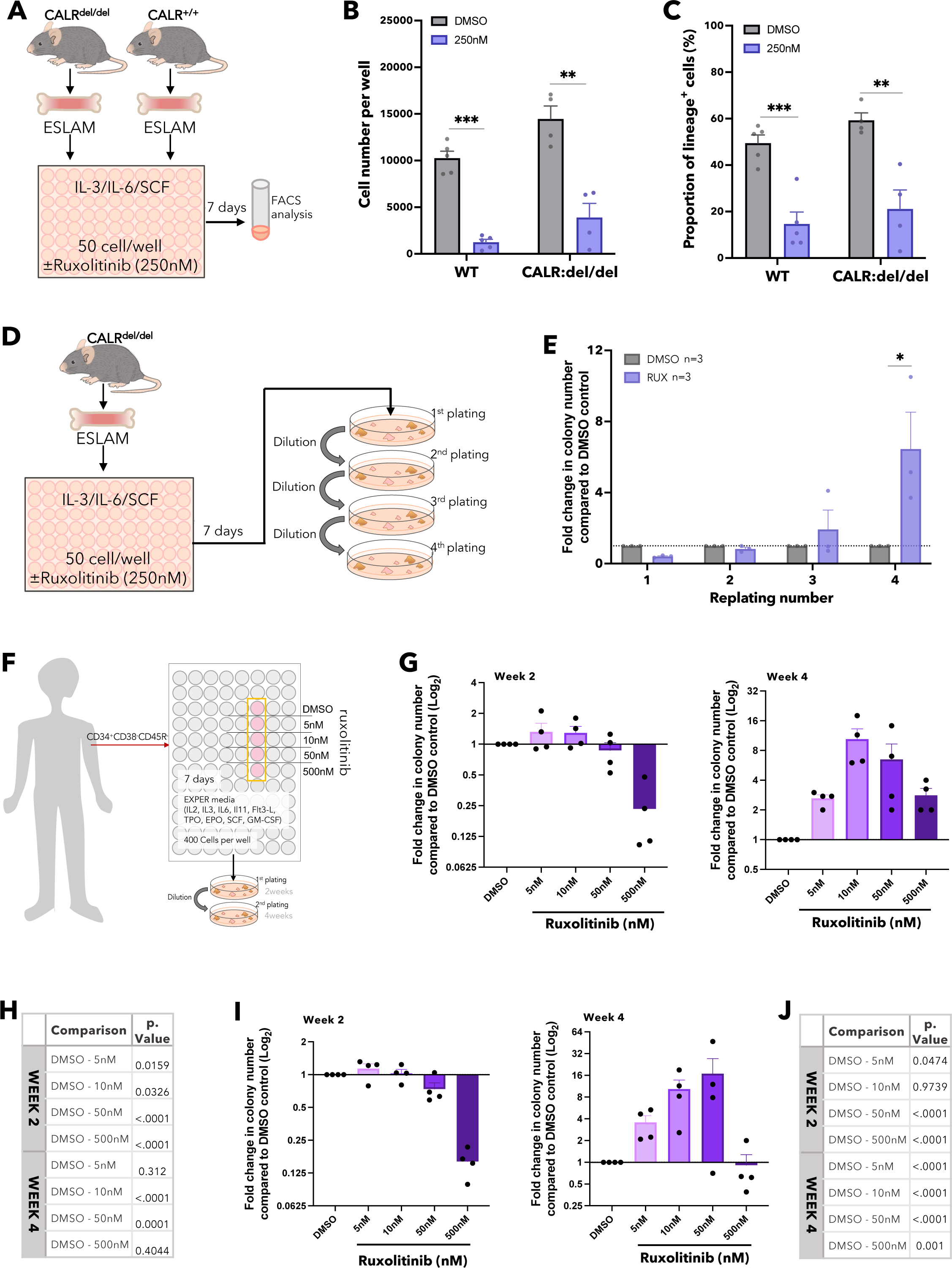
Ruxolitinib maintains murine and human myeloproliferative neoplasm HSPCs. (A) Schematic diagram showing *in vitro* functional assays of murine ESLAM HSCs treated with RUX or DMSO. ESLAM HSCs were FACS isolated from CALRdel/del (n=4) mutant mice and were then cultured for 7 days in IL-3/IL-6/SCF media^89^ with DMSO or 250nM of ruxolitinib before analysis by flow-cytometry. (B) Bar plots showing cell number per well in HSC derived cultures treated with vehicle or 250nM of ruxolitinib after 7 days (mean ± SEM). (C) Bar plots showing the proportion of cells expressing lineage positive markers (Ter119^+^/Ly6g^+^/CD11b^+^/B220^+^/CD3e^+^) after 7 days in culture with DMSO or 250nM of ruxolitinib (mean ± SEM). Asterisks indicate significant differences by Student’s t test (***, p<0.001; **, p<0.01; *, p<0.05). (D) Schematic diagram showing serial replating assays investigating the effect of Rux on ESLAM HSCs isolated from WT and CALRdel/del mutant mice. Sorted ESLAM HSCs were cultured for 7 days in IL-3/IL-6/SCF media^89^ with DMSO or 250nM of ruxolitinib and then subjected to serial colony replating assays. (E) Bar plots showing the fold change in number of colonies produced by HSC-derived cultures treated with vehicle or 250nM ruxolitinib for 7 days, normalised to the number of colonies produced by vehicle treated cultures at each week of replating. Results are from 2 independent experiments and are shown as mean ± SEM. Asterisks indicate significant differences by Mann-Whitney t test (*, p<0.05) (F) Schematic diagram showing MPP1-LTHSC (CD34^+^CD38^−^CD45RA^−^) cells were sorted from healthy human platelet apheresis donor cone samples, or myelofibrosis patient peripheral blood, into 96 well plates (400 cells/well) and cultured in high cytokine, serum-free medium (EXPER cytokine media)^71^ with scaled doses of ruxolitinib or vehicle control (DMSO). After 7 days the HSC derived cultures were plated in serial colony replating assays in methylcellulose. Healthy donors were all male and between 48-69 years of age. Myelofibrosis patient donors; three patients carried a JAK2 V617F mutation and were all male between the ages of 65-70, one donor carried a CALR-52bp deletion mutation and was a 70y/o female at the time of sample collection. (G) Bar plots showing the fold change in the number of colonies produced by HSPCs that were isolated from healthy donors and cultured for 7 days in the presence of ruxolitinib, normalised to the number of colonies produced by HSPCs cultured for 7 days with DMSO. Data shown as Log_2_(fold-change) from DMSO. Left showing fold change in colony numbers in the first round of colony formation (2 weeks in methylcellulose). Right showing fold change in colony numbers in the second round of colony formation (4 weeks in methylcellulose). Data from 4 healthy donors; each dot represents the mean fold change between technical replicates of a single donor. (H) Table showing the significance values (p.value) from estimated marginal (EM) means statistics derived from comparisons between DMSO and ruxolitinib conditions using a generalised mixed linear model applied to the raw colony counts used to generate 5G. (I) Bar plots showing the fold change in number of colonies produced by HSPCs that were isolated from patients with myelofibrosis and cultured for 7 days in the presence of ruxolitinib, normalised to the number of colonies produced by HSPCs cultured for 7 days with DMSO. Data shown as Log_2_(fold-change) from DMSO. Left showing fold change in colony numbers in the first round of colony formation (2 weeks in methylcellulose). Right showing fold change in colony numbers in the second round of colony formation (4 weeks in methylcellulose). Each dot represents the average fold-change from each of 4 patients with myelofibrosis and bars represent mean ± SEM. (J) Table showing the significance values (p.value) from EM means statistics derived from comparisons between DMSO and ruxolitinib conditions DMSO and ruxolitinib conditions using a generalised mixed linear model statistic applied to colony counts used to generate 5I.

To investigate whether ruxolitinib also maintained human HSPCs in *ex vivo* cultures, CD34^+^CD38^−^CD45RA^−^ progenitors were purified from apheresis cones derived from 4 platelet donors, grown in cytokine rich, serum-free culture conditions^71^ with or without ruxolitinib and their progeny assessed in serial colony replating assays (Fig. 6F). These human cell cultures did not contain albumin which binds ruxolitinib necessitating the use of lower ruxolitinib doses as previously described.^72^ After 2 weeks, ruxolitinib did not increase colony formation and even reduced colony output at the highest dose (500nM), but by 4 weeks it increased colony formation in all individuals at all doses tested, with 10nM and 50nM (similar to concentrations obtained in patients *in vivo*^73^ after accounting for albumin) showing the greatest benefit (Fig. 6G-6H, Fig. S6C-E and Table S5).

Ruxolitinib had a similar effect on HSCs derived from the peripheral blood of 4 patients with myelofibrosis with high WBC counts, none of whom had previously received ruxolitinib or interferon. Three patients were positive for the JAK2V617F mutation and one patient had a CALR-deletion mutation. After 2 weeks, ruxolitinib had little effect on colony output except at the highest dose, but at 4 weeks it substantially increased colony output in all 4 patients, with 10nM and 50nM concentrations showing the greatest benefit (Fig. 6I-J, Fig. S6F-H and Table S6).

Together, these data demonstrate that ruxolitinib maintained cultured murine myeloproliferative HSPCs, human normal HSPCs and human myeloproliferative HSPCs.

## Discussion

Our results demonstrate that STAT5 loss is accompanied not only by reduced HSC numbers but also a substantial HSC impairment associated with reduced cell cycle entry and increased differentiation. Prompted by this unusual phenotype, we show that uSTAT5 promotes maintenance and constrains differentiation of HSCs. Ruxolitinib, a JAK1/2 inhibitor in wide clinical use, increases uSTAT5 levels and enhances maintenance of WT and myeloproliferative HSCs from both mice and humans.

An intimate relationship between proliferation and differentiation has long been recognised in studies of HSC biology. Many genetic (e.g. ablation of CDKi^74–76^ or MEK1^77^) or environmental manipulations (e.g. infections or inflammation^78–81^) that induce HSC proliferation and functional exhaustion are associated with increased differentiation.^78–81^ In contrast, many of those that produce increased HSC quiescence are accompanied by reduced differentiation (e.g. Neo1 downregulation^82^ or Atad3a deletion^83^). However, we show here that highly purified STAT5-deficient HSCs display transcriptional evidence of reduced cell cycling together with functional evidence of reduced cell cycle entry, and yet are more prone to differentiation. Bunting and colleagues have previously reported that STAT5-deficient LSK or CD34^−^LSK HSCs displayed increased cell cycling.^26,51^ However, the frequencies of quiescent cells in their WT control populations were lower than those observed in ESLAM HSCs here (84% vs 91%), suggesting that cell populations gated for cell cycle analysis in the previous reports contained a higher frequency of more proliferative progenitors (presumably ST-HSC/MPP). The decreased frequency of primitive HSCs in STAT5^−/−^ mice likely resulted in a higher fraction of more proliferative ST-HSC/MPPs, thus increasing proliferation scores for populations containing such cells.

Our demonstration that STAT5 not only induces HSC proliferation but also represses HSC differentiation was reminiscent of previous results, which showed that uSTAT5 and pSTAT5 have separate transcriptional roles in megakaryocytic differentiation of multipotent HPC7 cells.^55^ We therefore explored the possibility that the functional consequences of STAT5 loss in HSCs might represent a compound phenotype involving loss of both uSTAT5 and pSTAT5 transcriptional programs. Three aspects of our studies are of particular note.

First, our results indicate that uSTAT5 constrains HSC differentiation (as shown by both loss-of-function and gain-of-function approaches) and also enhances HSC maintenance as assessed by serial replating and transplantation of STAT5-YF expressing cells. In the latter studies, STAT5-YF dramatically increased donor chimerism within the HSC compartment in both 1^0^ and 2^0^ recipients but reduced donor chimerism within whole bone marrow, indicating that STAT5-YF expressing HSCs are retained in the HSC compartment and are less likely to differentiate. Second, these functional changes reflected altered HSC transcriptional programs including signatures of reduced differentiation, increased quiescence and increased stemness as assessed by several different scoring systems. Third, our results highlight the need to take the signalling environment into account when interpreting the consequences of manipulating a STAT. Thus, using culture conditions that preclude significant STAT5 phosphorylation, the consequences of up or down-regulating STAT5 can be attributed to an effect on uSTAT5. By contrast when conditions are permissive for STAT5 phosphorylation it is difficult to disentangle effects attributable to uSTAT5 or pSTAT5.

It is interesting to consider our results in the light of data that HSCs can be expanded using culture conditions that include high TPO concentrations (100ng/ml).^84^ This observation contrasts with other reports showing that low TPO^85^ and low cytokine environments^86^ better maintain HSC function, and that injection of TPO or a TPO mimetic reduces HSC numbers and HSC function *in vivo*.^87^ Together these data indicate that the effect of TPO is complex and may be concentration and/or context dependent. TPO-driven HSC expansion may require other features of the “Wilkinson” expansion cultures (eg presence of PVA, absence of albumin, hypoxic incubation^84^).

Our results also have therapeutic implications. First they raise the possibility that ruxolitinib could be a useful strategy to enhance *ex vivo* maintenance of HSCs for gene therapies. Consistent with this concept JAK/STAT signalling is rapidly upregulated in cultured human HSCs and ruxolitinib improves maintenance of human HSPCs grown using gene therapy culture conditions (Johnson et al, in press). Second in patients with an MPN,^88,89^ JAK inhibitors have little if any effect on the level of the mutant clone.^50^ A protective effect of ruxolitinib on mutant HSCs, may contribute to the limited efficacy of JAK inhibitors. Third an accumulation of mutant HSCs poised to differentiate may also contribute to the JAK-inhibitor discontinuation syndrome, characterised by a rapid life-threatening MPN resurgence after JAK-inhibitor withdrawal.^88^ The cytokine environments of endogenous HSCs in their various niches remain poorly understood and so further studies will be needed to explore the HSC effects of ruxolitinib *in vivo*. However, our results raise the possibility that targeting uSTAT5 or total STAT5 activity may represent attractive therapeutic approaches for myeloid malignancies associated with JAK activation.

## Methods

### Mice

The wild-type C57BL/6 (CD45.2), C57BL/6.SJL (CD45.1) and F1 (CD45.1/CD45.2) mice, and CALR:del mutant mice^58^ in this study were used at 10-32 weeks of age. STAT5^fl/fl^mice^25^ were kindly gifted from Lothar Hennighausen and were crossed with Mx1Cre mice^59^ to generate STAT5^fl/fl^ with Cre (STAT5^fl/fl^Cre^+^) or without Cre (STAT5^fl/fl^Cre^−^). STAT5 deletion was induced with repeated injection with Polyinosinic:polycytidylic acid (Poly:IC). All mice were kept in pathogen free conditions and all procedures were performed according to UK Home Office regulations. Move some text from supplementary methods to here

### Poly:IC treatment

Poly:IC (γ-irradiated sodium salt, Sigma) was prepared in PBS and administered intraperitoneally (IP) to STAT5fl/flCre+/- or STAT5fl/flCre−/− mice (16mg/kg) up to 7 times over 21 days. Peripheral blood (PB) was collected via tail vein into EDTA (Ethylenediaminetetraacetic acid) coated microvette tubes (Sarstedt) at least 4 weeks after the final IP injection. Blood mononuclear cell DNA was extracted at room temperature (RT) for 5 minutes with diluted NaOH. After pH neutralization, a genomic PCR was set up to test the level of STAT5 recombination in peripheral blood.

### Bone marrow cell harvest

Tibia, femur and iliac crest bones from both hind legs were crushed in a mortar with FACS buffer (Calcium free and Magnesium free PBS; Gibco) containing 2%FBS (Gibco) and 5mM EDTA) and filtered through a 70-um strainer to obtain single cell suspensions. Equal volume of ammonium chloride solution (STEMCELL Technologies) was mixed with cell suspension gently and incubated for 10 min at 4°C to lyse red blood cells. Cells were then spun at 360 xg for 5 min, and bone marrow mononuclear cells (BMMNCs) were washed in 40mL FACS buffer, and then used for analyses.

### Flow cytometric analysis

Washed BMMNCs were resuspended in an appropriate volume for antibody staining. The BMMNC were incubated with the appropriate dilution of antibodies with fluorescent conjugates. Cells were washed in FACS buffer and then stained with 4’,6-diamidino-2-phenylindole (DAPI, Thermo Fisher) or 7AAD (Biolegend) for dead cell exclusion in FACS buffer. Stained cells were then analysed on a LSR Fortessa flow cytometer (BD Biosciences) equipped with FACSDiva Software. Data were analyzed using FlowJo (Tree Star).

The frequency of HSCs, progenitors and lineage cells in bone marrow was analyzed by flow cytometry: T cells, CD3e+; B cells, B220+; pro- and erythroblasts, defined by different levels of CD71 and Ter119 as in the figure legend; Monocytes/neutrophils, Ly6G+; Granulocytes/macrophages, Mac-1+; MK, CD41+CD42+; prog, Lin-Sca1-cKit+; megakaryocyte progenitors (MkP), Lin-Sca1-cKit+CD150+CD41+; GMP, Lin-Sca1-cKit+CD41-CD16/32+CD150-; PreCFU-e, Lin-Sca1-cKit+CD41-CD16/32-CD105+CD150+; CFU-e, Lin-Sca1-cKit+CD41-CD16/32-CD105+CD150-; PreMegE, Lin-Sca1-cKit+CD41-CD16/32-CD105-CD150+ and PreGM, Lin-Sca1-cKit+CD41-CD16/32-CD105-CD150-. Multipotent progenitor MPPs were defined as the following: MPP1 (Flk2−CD150+CD48−LSK), MPP2 (Flk2−CD150+CD48+LSK), MPP3 (Flk2−CD150−CD48+LSK) and MPP4 (Flk2+CD150−CD48+LSK). LT-HSC was defined as Flk2-CD34-Lin-Sca1+cKit+CD150+CD48- and ESLAM HSCs as CD45+EPCR+CD150+CD48-. CD45.1 and CD45.2 antibodies were used to distinguish donor, recipient and competitor cell populations in transplants. All antibodies were produced by Biolegend, except for anti-EPCR PE conjugated antibodies (STEMCELL Technologies), and anti-CD45.1 and anti-CD45.2 (BD Bioscience).

### FACS isolation of ESLAM HSCs

Single cell suspensions of BMMNCs were first lineage depleted using EasySep™ Mouse Hematopoietic Progenitor Cell Isolation Kit (STEMCELL Technologies). Lineage depleted cells were then stained with; EPCR (CD201) PE (clone RMEPCR1560, STEMCELL Tech), CD150 PE/Cy7 (clone TC15-12F12.2, BioLegend), CD45 FITC (clone 30-F1,1 BD Bioscience) and CD48 APC (clone HM48-1, Biolegend). ESLAM HSCs defined as CD45+EPCR+CD48-CD150+ as previously described,1 were isolated using a Becton Dickinson Influx sorter (BD Biosciences). Cells were sorted directly into culture dishes containing media or into Eppendorfs containing FACS buffer.

### Competitive transplantation assays

C57BL/6 (CD45.1) recipients were irradiated with 2 x 550 cGy. For competitive repopulation assays, ESLAM HSCs or GFP+ transduced HSCs (CD45.2+) were injected with 3-5 x 105 competitor bone marrow cells obtained from CD45.1/CD45.2 F1 mice into recipient mice. To evaluate qualitative differences of WT and STAT5−/− HSCs, 30 FACS isolated ESLAM HSCs were mixed with 3 x 105 competitor bone marrow cells obtained from CD45.1/CD45.2 F1 mice before being injected into CD45.1 recipient mice. Blood was analyzed every 28 days. At 6 months post transplantation, bone marrow cells from the primary recipient mice were assessed for donor-derived HSC chimerism using flow cytometry by staining bone marrow cells with LT-HSC (Lin-Sca1+cKit+CD150+CD48-CD34-Flk2-) as well as CD45.1 and CD45.2. Secondary transplantation was then performed using 5 x 106 nucleated bone marrow cells from primary recipients of ESLAM HSCs.

To evaluate the long-term functional capacity of uSTAT5B overexpressing HSCs, WT ESLAM HSCs (CD45.2+) were FACS sorted and transduced with lentivirus containing STAT5B-YF, or EV in SCF/IL-11 cultures2, and three days later 113 transduced donor cells were transplanted per recipient mouse (CD45.1+) with 3 x 105 competitor bone marrow cells obtained from CD45.1/CD45.2 F1 mice. Blood was analyzed every 28 days for 6 months before secondary transplants were set up by injecting 5 x 106 primary recipient bone marrow cells into irradiated secondary recipient mice. Blood was analyzed every 28 days for 6 months before harvesting. Donor-derived HSC chimerism was assessed using flow cytometry by staining bone marrow cells with ESLAM HSC panels as described above, as well as antibodies for CD45.1 and CD45.2.

To evaluate the effect of ruxolitinib treatments on HSC function, ESLAM HSCs (CD45.2+) were pre-cultured for 5 days in IL-3/IL-6/SCF culture conditions3 with vehicle or ruxolitinib before being injected with 3 x 105 competitor bone marrow cells obtained from CD45.1+ mice into irradiated recipient mice (CD45.1+/CD45.2+). Blood was analyzed every 28 days for 6 months before harvesting. Donor-derived HSC chimerism was assessed using flow cytometry by staining bone marrow cells with ESLAM HSC panels as described above, as well as antibodies for CD45.1 and CD45.2.

In all cases, peripheral blood was obtained and analyzed by flow cytometry for donor contribution to T cells (CD3e+), B cells (B220+), Monocytes/neutrophils (Ly6G+); Granulocytes/macrophages (Mac-1+). Cells were co-stained with antibodies for CD45.1 and CD45.2 to distinguish the donor origin of repopulated cells.

### ESLAM HSC *ex vivo* cultures

ESLAM HSCs were sorted from STAT5−/− or WT mouse bone marrow cultured as single cells in round-bottom 96-well plates (Corning) or in bulk cultures of >50 cells per well in flat bottom 96-well plates (Corning). Each well was preloaded with StemSpan SFEM (serum-free expansion medium, STEMCELL Technologies). Cell suspensions were then topped up with equal volume of StemSpan SFEM media containing cytokines as indicated below after cell sorting was completed. Cells were cultured in 37°C, 5% CO2. IL-11 and SCF cultures:2 Equal volume of medium was added to each well to a final concentration of 10% FBS (STEMCELL Technologies), 1% penicillin/streptomycin (Sigma-Aldrich), 1% L-glutamine (Sigma-Aldrich), stem cell factor (SCF; 250 ng/mL; STEMCELL Technologies), IL-11 (20ng/mL; STEMCELL technologies) and 0.1 mM β-mercaptoethanol (Gibco). IL-3/IL-6 and SCF cultures:3 Equal volume of medium was added to each well to a final concentration of 10% FBS (STEMCELL Technologies), 1% penicillin/streptomycin (Sigma-Aldrich), 1% L-glutamine (Sigma-Aldrich), stem cell factor (SCF; 250 ng/mL; STEMCELL Technologies), IL-3 (10 ng/mL; PeproTech), IL-6 (10 ng/mL; PeproTech) and 0.1 mM β-mercaptoethanol (Gibco).

### Lentiviral infection of ESLAM HSCs

1000-20,000 ESLAM HSCs from STAT5−/− or WT mice were sorted and split across wells of a 96 well plate (Corning). Cells were cultured in 50µL of IL-11 and SCF media2 for 2h before being supplemented with polybrene (Sigma-Aldrich) and up to 20uL of a lentivirus equipped with the gene of interest (STAT5B-WT or STAT5B-Y699F) driven by MND promoter in a pCCL-c-MNDUS-IRES-EGFP backbone. The following morning cells were replated in 12 well plates in 600ul to dilute the polybrene and virus. Three days after infection, cells were sorted for living (DAPI-, Thermo Fisher) and green fluorescent protein (GFP+) and used in in vitro assays and transplantation experiments.

### Smart-seq2 and 10x Genomics single-cell RNA analysis (scRNAseq)

Single ESLAM HSCs were FACS sorted from bone marrow mononuclear cells (BMMNCs) and processed using Smart-seq2 (accession number: GSE223366). Lineage^−^c-Kit^+^ (LK) cells were sorted from BMMNCs and processed using 10xChromium (10xGenomics, Pleasanton, CA; GSE223632). Sorted ESLAM HSCs were transduced with lentivirus containing EV, STAT5B-WT or STAT5B-YF. After a 5-day culture, GFP^+^DAPI^−^ cells were processed using 10xChromium (10xGenomics, Pleasanton, CA; GSE223680). Sorted ESLAM HSCs were cultured for 5 days with ruxolitinib or DMSO, which were then processed using 10xChromium (10xGenomics, Pleasanton, CA; GSE260462). All data were deposited in the National Center for Biotechnology Information (NCBI) Gene Expression Omnibus (GEO).

### 10X data projection and cell type annotation

For STAT5−/− and STAT5+/+ HSPC datasets, uSTAT5B overexpressing HSC culture datasets, and ruxolitinib treated HSC culture datasets; in order to acquire the cell type information for each cell, the cells from all the samples (done individually for each batch of each experiment) were projected onto 2 different pre-annotated reference landscapes: 1) Dahlin landscape consists of both LSK and LK populations13 and 2) Nestorowa landscape with mainly hematopoietic stem and progenitor populations.14 Highly variable genes from the reference landscape were used for the PCA calculation. New data was projected onto the reference landscapes using PCA projection. The 15 nearest neighbor cells from the reference landscape were calculated for each cell of the new data based on Euclidean distance and the cell type annotation was assigned as the most frequent cell type out of the 15 matched nearest neighbours. As the Nestorowa landscape has a more detailed annotation on immature populations whereas Dahlin landscape has a more detailed annotation on more mature populations, for the final cell type assignment, the Dahlin landscape was used as the key reference, from which the cells that were annotated as HSC or immature populations were replaced by the annotations from the Nestorowa landscape.

## Acknowledgements

We thank all the technicians in the Green, Laurenti and Göttgens labs for their valuable technical assistance; R. Schulte, and C. Cossetti at the CIMR Flow Cytometry Core Facility for assistance with cell sorting; S. Mendez-Ferrer, D. Prins, S. Loughran, J. Deuel, and T. Klampfl for valuable constructive discussions; Justyna Rak for facilitating approval of mouse experimental work; M. Paramor for help with 10x scRNAseq; B. Arnold, M. Feetenby, N. Lumley, H. Bloy, L. Smith and the all members of the AMB Animal Core Facility for excellent technical assistance, animal welfare and husbandry.

## Author Contributions

MJW and JL designed, conducted experiments and analysed data; XW and HPB performed bioinformatic analyses; QW and SJ helped with intracellular flow; HJP helped with blood phenotypic analysis of the STAT5 mouse model; GG helped with FACS sorting; NKW and SJK helped with scRNAseq; GSV, helped with mice studies; RH helped to analyse ChIPseq data; EL, PC, CJ, EC, GM, JB and NM helped with data interpretation and supervised experiments with human HSCs; TLH, DCP, RA and RS provided technical assistance; PC, EL and GM helped with human data statistical analysis; MJW, JL, BG and ARG wrote the manuscript; JL, BG and ARG supervised the study.

## Data and materials availability

All single-cell RNA-seq data are deposited and are publicly available. The STAT5^−/−^ and WT ESLAM Smartseq2 dataset is available under GSE223366. The STAT5^−/−^ and WT LK 10x dataset is available under GSE223632. The STAT5-YF and EV infected LT-HSC 10x dataset is available under GSE223680. The Ruxolitinib treated LT-HSC 10x dataset is available under GSE260462. Lentiviral STAT5 plasmids are available upon request.

## References

1. Orkin SH, Zon LI. Hematopoiesis: an evolving paradigm for stem cell biology. Cell. 2008;

2. Laurenti E, Göttgens B. From haematopoietic stem cells to complex differentiation landscapes. Nature. 2018;

3. Rossi L, Lin KK, Boles NC, et al. Less Is More: Unveiling the Functional Core of Hematopoietic Stem Cells through Knockout Mice. Cell Stem Cell. 2012;11(3):302– 317.

4. Pietras EM. Inflammation: a key regulator of hematopoietic stem cell fate in health and disease. Blood. 2017;130(15):1693–1698.

5. Collins A, Mitchell CA, Passegué E. Inflammatory signaling regulates hematopoietic stem and progenitor cell development and homeostasis. Journal of Experimental Medicine. 2021;218(7):e20201545.

6. Haas S, Trumpp A, Milsom MD. Causes and Consequences of Hematopoietic Stem Cell Heterogeneity. Cell Stem Cell. 2018;22(5):627–638.

7. Herrera SC, Bach EA. JAK/STAT signaling in stem cells and regeneration: from Drosophila to vertebrates. Development. 2019;146(2):dev167643.

8. Chen E, Staudt LM, Green AR. Janus Kinase Deregulation in Leukemia and Lymphoma. Immunity. 2012;36(4):529–541.

9. Gotthardt D, Trifinopoulos J, Sexl V, Putz EM. JAK/STAT Cytokine Signaling at the Crossroad of NK Cell Development and Maturation. Front Immunol. 2019;10:.

10. Owen KL, Brockwell NK, Parker BS. JAK-STAT Signaling: A Double-Edged Sword of Immune Regulation and Cancer Progression. Cancers (Basel). 2019;11(12):.

11. Brooks AJ, Putoczki T. JAK-STAT Signalling Pathway in Cancer. Cancers (Basel). 2020;12(7):.

12. Moriggl R, Topham DJ, Teglund S, et al. Stat5 Is Required for IL-2-Induced Cell Cycle Progression of Peripheral T Cells. Immunity. 1999;10(2):249–259.

13. Kieslinger M, Woldman I, Moriggl R, et al. Antiapoptotic activity of Stat5 required during terminal stages of myeloid differentiation. Genes Dev. 2000;14(2):232–244.

14. Dai X, Chen Y, Di L, et al. Stat5 Is Essential for Early B Cell Development but Not for B Cell Maturation and Function. The Journal of Immunology. 2007;179(2):1068 LP – 1079.

15. Socolovsky M, Fallon AEJ, Wang S, Brugnara C, Lodish HF. Fetal Anemia and Apoptosis of Red Cell Progenitors in Stat5a−/−5b−/− Mice: A Direct Role for Stat5 in Bcl-XL Induction. Cell. 1999;98(2):181–191.

16. Socolovsky M, Nam H, Fleming MD, et al. Ineffective erythropoiesis in Stat5a−/−5b−/− mice due to decreased survival of early erythroblasts. Blood. 2001;98(12):3261–3273.

17. Wang Y, Levy DE. Comparative evolutionary genomics of the STAT family of transcription factors. JAKSTAT. 2012;1(1):23–36.

18. Nakajima H, Liu X-W, Wynshaw-Boris A, et al. An Indirect Effect of Stat5a in IL-2– Induced Proliferation: A Critical Role for Stat5a in IL-2–Mediated IL-2 Receptor α Chain Induction. Immunity. 1997;7(5):691–701.

19. Imada K, Bloom ET, Nakajima H, et al. Stat5b Is Essential for Natural Killer Cell– mediated Proliferation and Cytolytic Activity. J Exp Med. 1998;188(11):2067 LP – 2074.

20. Udy GB, Towers RP, Snell RG, et al. Requirement of STAT5b for sexual dimorphism of body growth rates and liver gene expression. Proceedings of the National Academy of Sciences. 1997;94(14):7239 LP – 7244.

21. Teglund S, McKay C, Schuetz E, et al. Stat5a and Stat5b proteins have essential and nonessential, or redundant, roles in cytokine responses. Cell. 1998;

22. Yao Z, Cui Y, Watford WT, et al. Stat5a/b are essential for normal lymphoid development and differentiation. Proc Natl Acad Sci U S A. 2006;103(4):1000 LP – 1005.

23. Hoelbl A, Kovacic B, Kerenyi MA, et al. Clarifying the role of Stat5 in lymphoid development and Abelson-induced transformation. Blood. 2006;107(12):4898 LP – 4906.

24. Bunting KD, Bradley HL, Hawley TS, et al. Reduced lymphomyeloid repopulating activity from adult bone marrow and fetal liver of mice lacking expression of STAT5. Blood. 2002;99(2):479–487.

25. Cui Y, Riedlinger G, Miyoshi K, et al. Inactivation of Stat5 in Mouse Mammary Epithelium during Pregnancy Reveals Distinct Functions in Cell Proliferation, Survival, and Differentiation. Mol Cell Biol. 2004;24(18):8037 LP – 8047.

26. Wang Z, Medrzycki M, Bunting ST, Bunting KD. Stat5-deficient hematopoiesis is permissive for Myc-induced B-cell leukemogenesis. Oncotarget. 2015;

27. Wierenga ATJ, Vellenga E, Schuringa JJ. Maximal STAT5-Induced Proliferation and Self-Renewal at Intermediate STAT5 Activity Levels. Mol Cell Biol. 2008;

28. Schuringa JJ, Chung KY, Morrone G, Moore MAS. Constitutive Activation of STAT5A Promotes Human Hematopoietic Stem Cell Self-Renewal and Erythroid Differentiation. Journal of Experimental Medicine. 2004;200(5):623–635.

29. Watanabe S, Zeng R, Aoki Y, Itoh T, Arai K. Initiation of polyoma virus origin-dependent DNA replication through STAT5 activation by human granulocyte-macrophage colony-stimulating factor. Blood. 2001;97(5):1266–1273.

30. Gouilleux F, Wakao H, Mundt M, Groner B. Prolactin induces phosphorylation of Tyr694 of Stat5 (MGF), a prerequisite for DNA binding and induction of transcription. EMBO J. 1994;13(18):4361–4369–4369.

31. Brooks AJ, Dai W, O’Mara ML, et al. Mechanism of Activation of Protein Kinase JAK2 by the Growth Hormone Receptor. Science (1979). 2014;344(6185):1249783.

32. Mui AL, Wakao H, Kinoshita T, Kitamura T, Miyajima A. Suppression of interleukin-3-induced gene expression by a C-terminal truncated Stat5: role of Stat5 in proliferation. EMBO J. 1996;15(10):2425–2433.

33. Pallard C, Gouilleux F, Benit L, et al. Thrombopoietin activates a STAT5-like factor in hematopoietic cells. EMBO J. 1995;14(12):2847–2856.

34. Reich NC. STATs get their move on. JAKSTAT. 2013;2(4):e27080–e27080.

35. Croker BA, Kiu H, Nicholson SE. SOCS regulation of the JAK/STAT signalling pathway. Semin Cell Dev Biol. 2008;19(4):414–422.

36. Maurer B, Kollmann S, Pickem J, Hoelbl-Kovacic A, Sexl V. STAT5A and STAT5B-Twins with Different Personalities in Hematopoiesis and Leukemia. Cancers (Basel). 2019;11(11):1726.

37. Halim CE, Deng S, Ong MS, Yap CT. Involvement of STAT5 in Oncogenesis. Biomedicines. 2020;8(9):316.

38. Wingelhofer B, Neubauer HA, Valent P, et al. Implications of STAT3 and STAT5 signaling on gene regulation and chromatin remodeling in hematopoietic cancer. Leukemia. 2018;32(8):1713–1726.

39. Gu L, Vogiatzi P, Puhr M, et al. Stat5 promotes metastatic behavior of human prostate cancer cells in vitro and in vivo. Endocr Relat Cancer. 2010;17(2):481–493.

40. Moser C, Ruemmele P, Gehmert S, et al. STAT5b as molecular target in pancreatic cancer--inhibition of tumor growth, angiogenesis, and metastases. Neoplasia. 2012;14(10):915–925.

41. James C, Ugo V, le Couédic J-P, et al. A unique clonal JAK2 mutation leading to constitutive signalling causes polycythaemia vera. Nature. 2005;434(7037):1144–1148.

42. Baxter EJ, Scott LM, Campbell PJ, et al. Acquired mutation of the tyrosine kinase JAK2 in human myeloproliferative disorders. The Lancet. 2005;365(9464):1054–1061.

43. Nangalia J, Massie CE, Baxter EJ, et al. Somatic CALR Mutations in Myeloproliferative Neoplasms with Nonmutated JAK2. New England Journal of Medicine. 2013;369(25):2391–2405.

44. Levine RL, Wadleigh M, Cools J, et al. Activating mutation in the tyrosine kinase JAK2 in polycythemia vera, essential thrombocythemia, and myeloid metaplasia with myelofibrosis. Cancer Cell. 2005;7(4):387–397.

45. Klampfl T, Gisslinger H, Harutyunyan AS, et al. Somatic Mutations of Calreticulin in Myeloproliferative Neoplasms. New England Journal of Medicine. 2013;369(25):2379–2390.

46. Kralovics R, Passamonti F, Buser AS, et al. A Gain-of-Function Mutation of JAK2 in Myeloproliferative Disorders. New England Journal of Medicine. 2005;352(17):1779– 1790.

47. Verstovsek S, Mesa RA, Gotlib J, et al. A double-blind, placebo-controlled trial of ruxolitinib for myelofibrosis. New England Journal of Medicine. 2012;366(9):799–807.

48. Verstovsek S, Mesa RA, Gotlib J, et al. Efficacy, safety and survival with ruxolitinib in patients with myelofibrosis: results of a median 2-year follow-up of COMFORT-I. Haematologica. 2013;98(12):1865 LP – 1871.

49. Cervantes F, Vannucchi AM, Kiladjian J-J, et al. Three-year efficacy, safety, and survival findings from COMFORT-II, a phase 3 study comparing ruxolitinib with best available therapy for myelofibrosis. Blood. 2013;122(25):4047 LP – 4053.

50. Sureau L, Orvain C, Ianotto J-C, et al. Efficacy and tolerability of Janus kinase inhibitors in myelofibrosis: a systematic review and network meta-analysis. Blood Cancer J. 2021;11(7):135.

51. Wang Z, Li G, Tse W, Bunting KD. Conditional deletion of STAT5 in adult mouse hematopoietic stem cells causes loss of quiescence and permits efficient nonablative stem cell replacement. Blood. 2009;113(20):4856 LP – 4865.

52. Kollmann S, Grausenburger R, Klampfl T, et al. A STAT5B–CD9 axis determines self-renewal in hematopoietic and leukemic stem cells. Blood. 2021;138(23):2347–2359.

53. Bradley HL, Hawley TS, Bunting KD. Cell intrinsic defects in cytokine responsiveness of STAT5-deficient hematopoietic stem cells. Blood. 2002;100(12):3983 LP – 3989.

54. Li G, Wang Z, Zhang Y, et al. STAT5 requires the N-domain to maintain hematopoietic stem cell repopulating function and appropriate lymphoid-myeloid lineage output. Exp Hematol. 2007;35(11):1684–1694.

55. Park HJ, Li J, Hannah R, et al. Cytokine-induced megakaryocytic differentiation is regulated by genome-wide loss of a uSTAT transcriptional program. EMBO J. 2016;

56. Schepers H, van Gosliga D, Wierenga ATJ, et al. STAT5 is required for long-term maintenance of normal and leukemic human stem/progenitor cells. Blood. 2007;110(8):2880–2888.

57. Kato Y, Iwama A, Tadokoro Y, et al. Selective activation of STAT5 unveils its role in stem cell self-renewal in normal and leukemic hematopoiesis. J Exp Med. 2005;202(1):169 LP – 179.

58. Li J, Prins D, Park HJ, et al. Mutant calreticulin knockin mice develop thrombocytosis and myelofibrosis without a stem cell self-renewal advantage. Blood. 2018;131(6):649–661.

59. Kühn R, Schwenk F, Aguet M, Rajewsky K. Inducible Gene Targeting in Mice. Science (1979). 1995;269(5229):1427–1429.

60. Dahlin JS, Hamey FK, Pijuan-Sala B, et al. A single cell hematopoietic landscape resolves eight lineage trajectories and defects in Kit mutant mice. Blood. 2018;blood-2017-12-821413.

61. Nestorowa S, Hamey FK, Pijuan Sala B, et al. A single-cell resolution map of mouse hematopoietic stem and progenitor cell differentiation. Blood. 2016;128(8):e20–e31.

62. Kent DG, Dykstra BJ, Cheyne J, Ma E, Eaves CJ. Steel factor coordinately regulates the molecular signature and biologic function of hematopoietic stem cells. Blood. 2008;112(3):560–567.

63. Hamey FK, Göttgens B. Machine learning predicts putative hematopoietic stem cells within large single-cell transcriptomics data sets. Exp Hematol. 2019;78:11–20.

64. Wilson NK, Kent DG, Buettner F, et al. Combined Single-Cell Functional and Gene Expression Analysis Resolves Heterogeneity within Stem Cell Populations. Cell Stem Cell. 2015;

65. Cabezas-Wallscheid N, Buettner F, Sommerkamp P, et al. Vitamin A-Retinoic Acid Signaling Regulates Hematopoietic Stem Cell Dormancy. Cell. 2017;169(5):807–823.e19.

66. Pietras EM, Lakshminarasimhan R, Techner J-M, et al. Re-entry into quiescence protects hematopoietic stem cells from the killing effect of chronic exposure to type I interferons. Journal of Experimental Medicine. 2014;211(2):245–262.

67. Dykstra B, Kent D, Bowie M, et al. Long-Term Propagation of Distinct Hematopoietic Differentiation Programs In Vivo. Cell Stem Cell. 2007;1(2):218–229.

68. Alessandro M. Vannucchi, Peter A. W. te Boekhorst, Claire N. Harrison, et al. EXPAND, a dose-finding study of ruxolitinib in patients with myelofibrosis and low platelet counts: 48-week follow-up analysis. Haematologica. 2019;104(5):947–954.

69. Verstovsek S, Kantarjian H, Mesa RA, et al. Safety and efficacy of INCB018424, a JAK1 and JAK2 inhibitor, in myelofibrosis. New England Journal of Medicine. 2010;363(12):1117–1127.

70. Pecquet C, Chachoua I, Roy A, et al. Calreticulin mutants as oncogenic rogue chaperones for TpoR and traffic-defective pathogenic TpoR mutants. Blood. 2019;blood-2018-09-874578.

71. Belluschi S, Calderbank EF, Ciaurro V, et al. Myelo-lymphoid lineage restriction occurs in the human haematopoietic stem cell compartment before lymphoid-primed multipotent progenitors. Nat Commun. 2018;9(1):4100.

72. De Marinis E, Ceccherelli A, Quattrocchi A, et al. Ruxolitinib binding to human serum albumin: bioinformatics, biochemical and functional characterization in JAK2V617F+ cell models. Sci Rep. 2019;9(1):16379.

73. Shi JG, Chen X, McGee RF, et al. The Pharmacokinetics, Pharmacodynamics, and Safety of Orally Dosed INCB018424 Phosphate in Healthy Volunteers. The Journal of Clinical Pharmacology. 2011;51(12):1644–1654.

74. Zou P, Yoshihara H, Hosokawa K, et al. p57^Kip2^ and p27^Kip1^ Cooperate to Maintain Hematopoietic Stem Cell Quiescence through Interactions with Hsc70. Cell Stem Cell. 2011;9(3):247–261.

75. Matsumoto A, Takeishi S, Kanie T, et al. p57 Is Required for Quiescence and Maintenance of Adult Hematopoietic Stem Cells. Cell Stem Cell. 2011;9(3):262–271.

76. Cheng T, Rodrigues N, Shen H, et al. Hematopoietic Stem Cell Quiescence Maintained by p21cip1/waf1. Science (1979). 2000;287(5459):1804–1808.

77. Baumgartner C, Toifl S, Farlik M, et al. An ERK-Dependent Feedback Mechanism Prevents Hematopoietic Stem Cell Exhaustion. Cell Stem Cell. 2018;22(6):879–892.e6.

78. Essers MAG, Offner S, Blanco-Bose WE, et al. IFNα activates dormant haematopoietic stem cells in vivo. Nature. 2009;458:904.

79. Baldridge MT, King KY, Boles NC, Weksberg DC, Goodell MA. Quiescent haematopoietic stem cells are activated by IFN-γ in response to chronic infection. Nature. 2010;465:793.

80. Haas S, Hansson J, Klimmeck D, et al. Inflammation-Induced Emergency Megakaryopoiesis Driven by Hematopoietic Stem Cell-like Megakaryocyte Progenitors. Cell Stem Cell. 2015;17(4):422–434.

81. Bogeska R, Mikecin A-M, Kaschutnig P, et al. Inflammatory exposure drives long-lived impairment of hematopoietic stem cell self-renewal activity and accelerated aging. Cell Stem Cell. 2022;29(8):1273–1284.e8.

82. Renders S, Svendsen AF, Panten J, et al. Niche derived netrin-1 regulates hematopoietic stem cell dormancy via its receptor neogenin-1. Nat Commun. 2021;12(1):608.

83. Jin G, Xu C, Zhang X, et al. Atad3a suppresses Pink1-dependent mitophagy to maintain homeostasis of hematopoietic progenitor cells. Nat Immunol. 2018;19(1):29–40.

84. Wilkinson AC, Ishida R, Kikuchi M, et al. Long-term ex vivo haematopoietic-stem-cell expansion allows nonconditioned transplantation. Nature. 2019;571(7763):117–121.

85. Kobayashi H, Morikawa T, Okinaga A, et al. Environmental Optimization Enables Maintenance of Quiescent Hematopoietic Stem Cells Ex Vivo. Cell Rep. 2019;28(1):145–158.e9.

86. Oedekoven CA, Belmonte M, Bode D, et al. Hematopoietic stem cells retain functional potential and molecular identity in hibernation cultures. Stem Cell Reports. 2021;16(6):1614–1628.

87. Nakamura-Ishizu A, Matsumura T, Stumpf PS, et al. Thrombopoietin Metabolically Primes Hematopoietic Stem Cells to Megakaryocyte-Lineage Differentiation. Cell Rep. 2018;25(7):1772–1785.e6.

88. Palandri F, Palumbo GA, Elli EM, et al. Ruxolitinib discontinuation syndrome: incidence, risk factors, and management in 251 patients with myelofibrosis. Blood Cancer J. 2021;11(1):4.

89. Daniel P, Jung PH, Sam W, et al. The stem/progenitor landscape is reshaped in a mouse model of essential thrombocythemia and causes excess megakaryocyte production. Sci Adv. 2022;6(48):eabd3139.

