## Supplemental Information for "Maintenance of haematopoietic stem cells by JAK inhibition and increased tyrosine-unphosphorylated STAT5"

**Supplemental data for Williams et al.**

- Supplemental methods
- Supplemental Figures SF1-SF6
- Supplemental Tables ST1-ST6
- References

#### Supplemental Methods

##### Single cell *in vitro* division kinetic assays

Single STAT5$^{-/-}$ or WT ESLAM HSCs were sorted at single cell into round bottom 96-well plates (Corning). Cells were cultured in 100μL IL-11 and SCF culture media[2]. Cell counts were scored daily under microscope, and cell cycle kinetics was determined as described (Li et al 2018) for the first, second, third and fourth division based visually scored cell numbers. First division (≥2 cells per well), second division (≥3 cells per well), third division (≥5 cells per well), or fourth division (≥8 cells per well).

##### Bulk ESLAM HSC *in vitro* expansion and differentiation analysis

50 STAT5$^{-/-}$ or WT ESLAM HSCs were sorted into flat-bottom 96-well plates (Corning). For STAT5-deficient HSC expansion and differentiation analysis at days 4, 5, 6 and 10, cells were cultured in IL-11 and SCF conditions.[2] For ruxolitinib treated HSC differentiation and expansion analysis, cells were placed in IL-3, IL-6 and SCF cultures conditions[3] for 2h and then ruxolitinib was applied in equal volume to final concentrations indicated in figures and cultured for 7 days. Cells were harvested at indicated time points and were then washed with FACS buffer and resuspended in an appropriate volume of FACS buffer for antibody staining. Cells were stained with fluorescently labelled antibodies against: Ly-6G BV785 (Biolgend), CD11b AF700 (Biolegend), Ter119 PE (Biolegend), B220 FITC (Biolegend), CD3e FITC (Biolegend), CD19 FITC (Biolegend), NK1.1 APC (Biolegend), c-Kit APC/Cy7 (Biolegend), Sca1 BV605 (Biolegend). Alternatively cells were stained with Lineage cocktail AF700 (Biolegend), which contained anti

Ter119, CD3e, B220 Ly6G and CD11b antibodies. Cells were washed in FACS buffer and then stained with 4',6-diamidino-2-phenylindole (DAPI, Thermo Fisher) or 7AAD (Biolegend) for dead cell exclusion in FACS buffer. Stained cells were then analyzed on LSRFortessa flow cytometer (BD Biosciences) equipped with FACSDiva Software. Data were analyzed using FlowJo (Tree Star). Accurate cell counts were acquired by applying a defined number of fluorescent beads (Trucount Control Beads, BD Biosciences) to each well. Each sample was back-calculated to the proportion of the total that were run through the cytometer.

**Analysis of STAT5B-YF transduced ESLAM HSC single cell cultures**

Transduced ($GFP^+$) $STAT5^{-/-}$ or WT ESLAM HSCs were sorted 3 days post infection at single cell per well in round-bottom 96-well plates (Corning) and cultured in IL-11 and SCF media[2] for a further 7 days. Cells were harvested and washed with FACS buffer and resuspended in an appropriate volume of FACS buffer for antibody staining and analyzed for lineage positive cell frequency and cell number. Stained cells were then analyzed on LSRFortessa flow cytometer (BD Biosciences) equipped with FACSDiva Software. Data were analyzed using FlowJo (Tree Star). Accurate cell counts were acquired by applying a defined number of fluorescent beads (Trucount Control Beads, BD Biosciences) to each well. Each sample was back-calculated to the proportion of the total that were run through the cytometer.

**Lentivirus production**

Lentiviral production was performed with HEK Lenti-X 293T cells (Takara) in low glucose Dulbecco's Modified Eagle's Medium with GlutaMAX and sodium pyruvate

(DMEM, Thermo Fisher), supplemented with 10% heat-inactivated FBS (Sigma-Aldrich), 1% penicillin/streptomycin (Sigma-Aldrich). HEK Lenti-X 293T cells were split 24 hours before production protocol began.

HEK Lenti-X 293T cells were plated at 9 million cells in 20mL media in a 15cm dish (Corning) and cultured for 24h. Two hours before transfection with the plasmids, media was replaced with 18mL fresh media. 2ml of DMEM containing 3$^{rd}$ generation lentiviral packaging plasmids (12µg, Takara), Lipofectamine (3% f.c. Thermo Fisher), and plasmid of interest (12µg, pCCL-c-MNDUS-IRES-EGFP containing EV, STAT5B-WT or STAT5B-Y699F, were added to dishes while swirling. Plates were incubated over-night and media was changed the following morning. 48h after transfection, media was collected and filtered through 0.45µm syringe filters (Falcon). Virus was concentrated with Lenti-X concentrator (Takara) according to manufacturer's instruction. 20µl aliquots were snap frozen and then stored at -80°C.

**Mouse HSPC serial colony replating assays**

For all serial colony replating assays, cells were seeded into M3434 methylcellulose (STEMCELL Tech) without antibiotic in STEMvision 6-well plates and were allowed to grow for 1 week before imaging the plate on STEMvision (STEMCELL Technologies). The entire well contents of each condition was harvested and a fraction of cells were seeded into M3434 methycellulose and were allowed to grow for 1 week before imaging. This protocol was performed until colonies were no longer seen across all conditions. Colony Forming Unit (CFU) was defined here as the number of colonies observed, multiplied by the dilution factor (which gets greater with each passing week), divided by the starting number of HSC cells.

For uSTAT5B analysis; transduced STAT5$^{-/-}$ or WT ESLAM HSCs (DAPI$^{-}$GFP$^{+}$) were sorted 3 days post infection and were seeded at 75 cells per well and split over two duplicate wells. For ruxolitinib analysis; 50 WT ESLAM HSCs were placed into flat bottom 96 well plates (Corning) in SCF, IL-3 and IL-6 cultures.[3] After 2 hours, cultures were treated with DMSO or ruxolitinib (250nM and 1000nM) for 7 days. Diluted cultured cells were then plated in methylcellulose.

**Cell cycle analysis of ESLAM HSCs**

Bone marrow cells from STAT5$^{-/-}$ and WT control mice were lineage depleted using EasySep™ Mouse Hematopoietic Progenitor Cell Isolation Kit (STEMCELL tech). The cells were then stained for 45 minutes with CD45 BV785 (Biolegend), EPCR PE (STEMCELL Technologies), CD150 PE/Cy7 (Biolegend), CD48 BV605 (Biolegend) and Zombi Nir (Biolegend) to eliminate dead cells. The stained cells were washed with FACS buffer (PBS/2% FBS), then fixed and permeabilized using Cytofix/Cytoperm™ Fixation/Permeablization kit (BD Biosciences). The cells were then stained with Ki-67 FITC (Biolegend) on ice for 1 hr and then with DAPI (Thermo Fisher) for 1 hr at RT. Flow cytometric analysis was carried out at a low flow rate using LSRFortessa flow cytometer (BD Biosciences).

**Intracellular flow cytometry analysis of cultured ESLAM HSCs**

ESLAM HSCs from STAT5$^{-/-}$ and WT control mice were sorted and placed into StemSpan SFEM media (STEMCELL technologies) for 1h (cytokine starvation).

For intracellular flow analysis of STAT proteins, ruxolitinib or DMSO was added to StemSpan SFEM during the 1h starvation period. Cells were then stimulated with IL-11/SCF cultures,[2] cytokines present in M3434, or IL-3/IL-6/SCF cultures[3] for 30minutes. Cells were then fixed and permeabilized using Cytofix buffer and Perm Buffer III (BD Biosciences). The cells were then stained with anti-pY694/Y699 STAT5 PE antibody (Clone 47/Stat5(pY694), BD Bioscience) on ice for 1 hr. In Ruxolitinib experiments, cells were additionally stained with antibodies for pY701 STAT1 AF488 (Clone 4a, BD Bioscience), pY705 STAT3 AF647 (Clone 4/P-STAT3, BD Bioscience), and total STAT5 (Clone D2O6Y, Cell Signalling Technology) followed by staining with a PE anti-rabbit secondary antibody. STAT5-deficient cells were used a negative control for total-STAT5 antibody staining. Flow cytometric analysis was carried out on a LSR Fortessa flow cytometer (BD Biosciences) equipped with FACSDiva Software. Data were analyzed using FlowJo (Tree Star).

**Intracellular flow cytometry analysis of cultured HSPCs with STAT5-YF expression.**

HSPCs (LSK cells) from WT mice were sorted and placed into IL-11/SCF cultures,[2] for two hours before lentiviral transduction (see above). 3 days later, GFP+ cells were FACS isolated and placed into placed into StemSpan SFEM media (STEMCELL technologies) for 1h (cytokine starvation). Cells were then stimulated with IL-11/SCF cultures.[2] Cells were then fixed and permeabilized using Cytofix buffer and Perm Buffer III (BD Biosciences). The cells were then stained with anti-pY694/Y699 STAT5 PE antibody (Clone 47/Stat5(pY694), BD Bioscience) or total STAT5 (Clone D2O6Y, Cell Signalling Technology) followed by staining with a PE anti-rabbit secondary antibody on ice. Flow cytometric analysis was carried out on a LSR Fortessa flow cytometer (BD Biosciences) equipped with FACSDiva Software. Data were analyzed using FlowJo (Tree Star).

**Apoptosis analysis of lineage depleted bone marrow**

Bone marrow cell suspensions were mixed with equal volume of ammonium chloride (StemCell Technologies) and incubated on ice for 10 minutes to lyse red blood cells. Cells were then washed in 40ml of FACS buffer (PBS/2% FBS 5mM EDTA) and then lineage depleted using EasySep™ Mouse Hematopoietic Progenitor Cell Isolation Kit (STEMCELL technologies). Lineage depleted cells were cultured in 1.5mL IL-3, IL-6 and SCF culture in 6 well plates. After 2h in culture, 1.5mL media containing ruxolitinib (at 2x concentration of the indicated doses ) was added to wells and cultured over-night. Cells were then washed with FACS buffer and resuspended in an appropriate volume of FACS buffer for antibody staining. Cells were stained with c-Kit APC/CY7 (Biolegend) and Lin cocktail AF700 (Biolegend) for 45 minutes. The stained cells were washed with FACS buffer, then stained with 5ul Annexin V FITC (Biolegend) in 100µL 1x Annexin binding buffer at room temperature in dark for 15 minutes. DAPI (Thermo Fisher) was then added to the samples. Flow cytometric analysis was carried out using LSRFortessa flow cytometer (BD).

**Clone survival analysis of single HSC in *in vitro* culture**

Single WT HSCs or $GFP^+DAPI^-$ transduced cells were seeded into round bottom 96 well plates (Corning) in IL-3, IL-6, and SCF conditions[3] for ruxolitinib treatment analysis of clone viability or in IL-11 and SCF conditions[2] for uSTAT5B overexpression analysis of clone viability. Ruxolitinib was applied after a 2h rest period in culture as above. Wells were studied under a microscope after 5 days. The number of wells carrying 1 or more cells were marked as surviving clones. The frequency of surviving clones from the total clones plates was calculated.

**qPCR analysis**

WT HSCs were cultured in SCF IL-3 and IL-6 conditions[3] with ruxolitinib or DMSO for 2h before being collected and washed. RNA was extracted using PicoPure RNA isolation kit (Thermo Fisher) as per manufacturer's instruction. First strand cDNA synthesis was performed using SuperScript III First-Strand Synthesis System for RT-PCR (Thermo Fisher), using Random Hexamers, as per the manufacturer's guidelines. QPCR was carried out using PowerTrack™ SYBR Green Master Mix (Thermo Fisher) and the primers below. qPCR assays were run and analysed using Stratagene Mx3000P qPCR System. The expression of individual genes relative to a housekeeping gene (Gapdh) was calculated as R = 2-(CT target – CT housekeeping).

**Cish:**

Forward primer AGGGATCTTGTCCTTTGCTGG
Reverse primer AGGGTCTAGCACCTTCGGTT

**Pim1:**
Forward primer CAAGATCAACTCCCTGGCCC
Reverse primer TTAATGGCCACCGGCAAGTT

**HEL and BaF3 cell culture and western blot**

All cells were cultured in 37°C, 5% $CO_2$. HEL cells carrying a $JAK2^{V617F}$ mutation were cultured in RPMI-1640 media (Thermo Fisher) with 10% FBS (Gibco) and 1% penicillin/streptomycin (Sigma-Aldrich). Ba/F3 cells that overexpress c-MPL and mutant calreticulin (CALR:del) were cultured in RPMI-1640 media (Thermo Fisher) with 10% FBS (Gibco) and 1% penicillin/streptomycin (Sigma-Aldrich) with 25ng/mL of THPO (Thermo Fisher).

1 x $10^6$ cells were plated into 6cm dishes. 24h later cells were treated for 2h with ruxolitinib at 1000nM, 250nM or DMSO. Cells were washed with cold PBS (Gibco) and harvested in dishes on ice with RIPA buffer (Thermo Fisher) supplemented with protease and phosphatase inhibitors (Merck). Blots were stained with anti-STAT5 (D2O6Y, Cell Signaling), anti-pY694-STAT5 (ab32364, Abcam) and anti-tubulin (2144, Cell Signalling) in Tris buffered saline (TBS, Thermo Fisher) with 0.1% Tween and 5% BSA. Blots were stained with secondary HRP linked antibodies (Thermo Fisher) in TBS with 0.1% Tween and 3% milk powder (Marvel).

**Statistical Analysis**

For all comparisons of datasets where there was ≥3 replicate samples tested, an unpaired Student's t-test was tested in Graphpad (Prism) where all bar charts, dot plots and line graphs were plotted. The level of significance was indicated with asterisks. For comparing between conditions across multiple time points ($STAT5^{-/-}$ and WT cohort blood analysis, or $STAT5^{-/-}$ and WT competitive transplants blood repopulation) an ANOVA was performed and the column factor p value was indicated.

**Generation of $STAT5^{-/-}$ and WT ESLAM cells Smart-seq2 scRNAseq dataset**

Bone marrow cells were harvested from $STAT5^{-/-}$ and WT control mice and single ESLAM HSCs were FACS sorted after lineage depletion, as described above, and

RNA libraries were made according to manufacturer's protocol (Illumina) processed by as described for Smart-seq2.

**Preprocessing Smart-seq2 data**

The reads resulting from the SmartSeq2 experiments were mapped against Ensembl genes (release 81)[4] using GSNAP (version 2015-09-29)[5] and quantified using HTSeq (version 0.6.0).[6] Filtering Quality Control (QC) steps were applied where nuclear genes had to be at least 20% of the mapping reads and cells with less than 50,000 reads mapping to them were rejected. In addition, the maximum allowed fraction of cells mapping to mitochondrial genes was set at 20%. The levels of technical variance were estimated using the ERCC spike-ins as described by Brennecke *et al.* (2013)[7] with highly variable genes (HVGs) being defined as having the squared coefficient of variation exceeding technical noise. The resulting dataset was also transformed by applying the remove batch effect method implemented within the R limma package.[8]

**Additional processing and filtering Smart-seq2 data**

Further processing of the dataset was performed with the python package Scanpy.[9] The dataset was subsequently log-transformed, scaled and using the HVGs as input a PCA reduction for the top 50 components was computed. A diffusion map embedding was also calculated for the top 15 diffusion components. While considering the geometric mean of Stat5a/Stat5b expression 23 HSCs from STAT5 deficient mice were excluded from further analysis. These excluded cells corresponded to the top 20% *Stat5a* and *Stat5b* expressing HSCs from the STAT5 deficient mice.

**Analysis of differential expression genes of the Smart-seq2 dataset**

Differential expression on the Smart-seq2 sequencing data was performed using the R package DESeq2.[10] The input expression matrices were filtered to include only genes with average expression above 1 (resulting in 14,080 genes on the filtered matrix). Differential expression results were selected for a significance level of 0.01. Visualization of results was done via volcano plot generated with the Enhanced Volcano R package.[11]

**Gene set enrichment analysis of the Smart-seq2 data**

The Wald statistic from DESeq2 result was used to generate pre-ranked gene lists. These gene lists were then used for Gene set enrichment analysis (GSEA) with the GSEA software (v3.0) pre-ranked mode. Enrichment was tested for Hallmark and KEGG curated genesets of the MSigDB database v7.0.[12]

**Generation of 10x scRNAseq dataset of STAT5$^{-/-}$ and WT**

Bone marrow cells were harvested from STAT5KO and WT control mice and lineage negative, c-Kit$^{+}$ (LK) cells were FACS sorted and processed according to the manufacturer's protocol for the 10x Chromium™ Single Cell 3' kit (v2)(10x Genomics, Pleasanton, CA). There were six samples in total, including 2 replicates for 3 months WT samples, 2 replicates for 3 months Stat5KO samples and 1 sample each for 12 months WT and Stat5KO, respectively. The prepared libraries were sequenced on a Novaseq 6000. The sequenced reads were processed with Cellranger (version 2.1.1) and aligned to the 10X Genomics built mouse mm10 reference (version 1.2.0).

**STAT5$^{-/-}$ and WT LK 10X data pre-processing**

The Scrublet package was used to estimate doublets and 449 were selected and removed as such. The cells were further filtered based on the percentage of mitochondrial UMI counts being less than 10% per cell total and cells expressing at least 500 genes. Genes were remained if they were expressed in at least in 3 cells. The filtered dataset was subsequently normalized to the median UMI counts per cell across the two processed libraries and log-transformed.

**Downstream analysis of the STAT5$^{-/-}$ and WT LK 10X scRNAseq dataset**

After the preprocessing step, there were 30728 cells past quality control (3m WT S1: 5708; 3m WT S2: 4943; 3m Stat5KO S1: 4469; 3m Stat5KO S2: 4596; 12m WT: 5239 and 12m Stat5KO: 5773). The following downstream analysis was performed using the Scanpy package in Python. Cell cycle phases were estimated using the score_genes_cell_cycle function in Scanpy with the default 97 cell cycle genes defined in Tirosh et al, 2015. Highly variable genes were then computed with parameters *min_mean*=0.02, *max_mean*=3, *min_disp*=0.3 within each batch separately and merged. The effect of cell cycle phases, number of genes, number of total counts and percentage of mitochondrial genes were removed using the regress_out function in Scanpy. The dataset was scaled prior to the principle component analysis (PCA) calculation. Uniform Manifold Approximation and Projection (UMAP), a dimension reduction technique was used to visualise the data using the top 50 PCs.

**Generation of 10X scRNAseq dataset of STAT5B-YF ESLAM HSC *in vitro* culture**

ESLAM HSCs were sorted as $CD45^+EPCR^+CD150^+CD48^-$ and transduced with lentivirus containing STAT5B-YF or EV, respectively, in SCF and IL-11 cultures. Five days after infection, $DAPI^-$ $GFP^+$ were sorted for 10X scRNAseq. There were 2 replicates for each condition. The cells were processed with a Chromium Controller according to the Chromium NextGEM Single Cell 3' GeneExpression v3.1 protocol. The libraries were pooled and sequenced on a Novaseq 6000.

**Pre-processing of the 10X scRNAseq dataset of STAT5B-YF or EV transduced ESLAM HSC *in vitro* culture**

The scrublet package in Python was used to estimate doublets and cells estimated to potentially be doublets were removed. In addition, cells with less than 1000 genes, or more than 2e5 UMI counts or more than 5% mitochondrial gene counts were further eliminated to get best quality cells for downstream analysis.

**Downstream analysis of the 10X scRNAseq dataset of STAT5B-YF ESLAM HSC *in vitro* culture**

After the pre-processing step, there were 21649 cells remaining (629; YF B1: 494; EV B1: 629; YF B2: 3945 and EV B2: 10149). Highly variable genes were then computed with parameters *min_mean*=0.02, *max_mean*=3, *min_disp*=0.5 within each batch separately and merged. The effect of cell cycle phases, number of genes, number of total counts and percentage of mitochondrial genes were removed using the regress_out function in Scanpy. The dataset was scaled prior to the principle component analysis (PCA) calculation. Uniform Manifold

Approximation and Projection (UMAP), a dimension reduction technique was used to visualise the data using the top 50 PCs.

**Generation of 10x scRNAseq dataset of Ruxolitinib treated ESLAM HSC culture**

ESLAM HSCs were sorted and cultured in the medium that contains SCF, IL3 and IL6 to promote STAT5 phosphorylation with DMSO or ruxolitinib (250nM) for 5 days. After that, DAPI$^-$ cells were sorted for scRNAseq. There were 2 replicates for each condition. The cells were processed with a Chromium Controller according to the Chromium NextGEM Single Cell 3' GeneExpression v3.1 protocol. The libraries were pooled and sequenced on a Novaseq 6000. The sequenced reads were processed with the cellranger pipeline (version 3.1.0) and aligned to the genome reference 10X Genomics-builqd mouse mm10 (version 3.0.0).

**Pre-processing of the ESLAM HSC culture treated with DMSO or Ruxolitinib**

The Scanpy (v1.9.6) package in Python (v3.9.18) was used for data pre-processing. The scrublet package (v0.2.3) in Python was applired to estimate doublets and cells estimated to potentially be doublets were removed. In addition, cells with less than 500 genes, or more than 2e5 UMI counts or more than 5% mitochondrial gene counts were further eliminated to get best quality cells for downstream analysis.

**Downstream analysis of the 10x scRNAseq dataset of Ruxolitinib treated ESLAM HSC culture**

After the pre-processing step, there were 19565 cells remaining (DMSO_rep1: 11428, DMSO_rep2:4972, Rux_rep1:1123, Rux_rep2:2042). The cells were normalized to a total count of 1e4. Highly variable genes were then computed with parameters min_mean=0.02, max_mean=3, min_disp=0.5 within each batch separately and merged. The effect of cell cycle phases, number of genes, number of total counts and percentage of mitochondrial genes were removed using the regress_out function in Scanpy. Cell cycle phases were estimated using the score_genes_cell_cycle function in Scanpy. After that, the cell cycling effect was further eliminated using the regress_out function. Harmony (v1.2.3360) was then applied to correct potential batch effects between samples. The dataset was scaled prior to the principle component analysis (PCA) calculation. Umap was used to visualise the data using the top 50 PCs. Cell types were annotated using the published Nestorowa [PMID:27365425] and Dahlin [PMID:29588278] landscapes. Leiden clustering with resolution=0.8 defined 15 clusters, of which cluster 10-12 were annotated as non-HPSC cells and hence were removed for further analysis. Differential expression analysis was carried out for each cell type between Rux-treated and DMSO-control using the edgeR package (v4.0.5) in R (v4.3.1). Pre-filter on genes was done using min.count=1, min.total.count=1, large.n=10 and min.prop=0.3. All the plots were created using ggplot2 (v3.4.4) in R.

**Visualizations**

All violin plots were done using the ggplot2 package in R. Mean difference between groups were tested using t test.

**Human samples from healthy donors**

Non mobilized peripheral blood samples were collected from leukocyte cones (a by-product of platelet apheresis) with informed consent by NHS Blood and

Transfusion (NHSBT) Centres and sourced by the Cambridge Blood and Stem Cell Biobank. All experimental work was performed in accordance with regulated procedures approved under the 07/MRE05/44 and 18/EE/0199 REC research studies.

**Human samples from myelofibrosis patients**

Non mobilized peripheral blood samples were collected from the peripheral blood of myelofibrosis (MF) patients, all with informed consent by NHS Blood and Transfusion (NHSBT) Centres and sourced by the Cambridge Blood and Stem Cell Biobank. All experimental work was performed in accordance with regulated procedures approved under the 18/EE/0199 REC research studies. All patients (n=4) had white blood cell counts greater than $10x10^9$/L and had not received ruxolitinib or interferon as part of their medical treatment prior to sample collection. 3 donors were diagnosed with JAK2V617F mutant MF and one patient was diagnosed with CALR-Del mutant MF.

**Human serial colony replating**

Non mobilized peripheral blood HSC pool cells ($CD34^+CD38^-CD45RA^-$) were isolated from healthy donors or MF patients and seeded at 400 cells per well in 96 well plates and cultured in conditioned media termed "MEM" media.[15] MEM media: StemPro base media supplemented with Nutrients (0.035%), Pen/Strep (1%), L-Glu (1%), human LDL (50ng/ml), SCF (100ng/ml), Flt-3L (20ng/ml), TPO (100ng/ml), EPO (3 units/ml), IL-6 (50ng/ml), IL-3 (10ng/ml), GM-CSF (20ng/ml), IL-11 (50ng/ml), IL-2 (10ng/ml) and IL-7 (40ng/ml). All cytokines from Miltenyi Biotech except EPO (Janssen). Two hours , RUX (5nM, 10nM, 50nM or 500nM,

Selleckchem) or DMSO (Sigma) was added to the media. After 5 days of culture, cells were harvested and washed with FACS buffer (PBS + 3% FBS). Cells from each condition were split into two dilutions: either containing 20% or 80% of cells. These two dilutions were placed into H4034 Methocult medium (Stemcell Technologies) supplemented with 10ng/mL Flt3-L (PeproTech), 10ng/mL IL-6 (PeproTech) and 10U/mL Penicillin-Streptomycin (Thermo Fisher Scientific) and split into duplicate wells (2 x 10% cell suspension and 2 x 40% cell suspension) of a 6 well plate. After 14 days the number of colonies in each condition was counted using the Stemvision analyser (Stemcell Technologies) (primary plating analysis). After counting, the duplicate well contents were harvested and washed with FACS buffer (3% FBS in PBS). The harvested cells were again split into two dilutions (either containing 20% or 80% of cells), and placed into H4034 Methocult (Stemcell Technologies) with added cytokines and Penicillin-Streptomycin, and were then split into duplicate wells. After a further 14 days the colonies were imaged and counted (secondary plating analysis).

### Supplemental Figures

#### Supplemental Figure 1: STAT5 loss results in defective HSC function

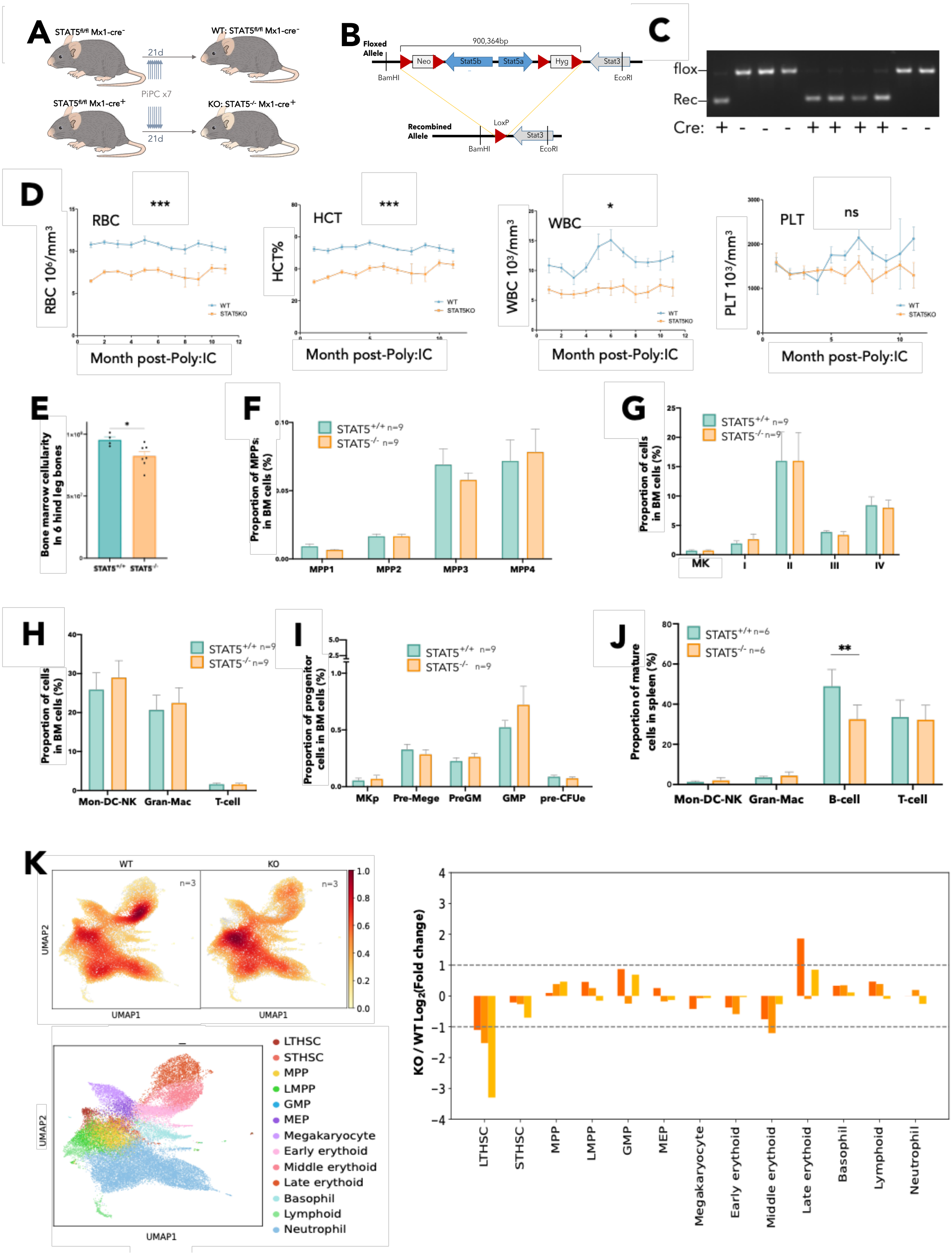

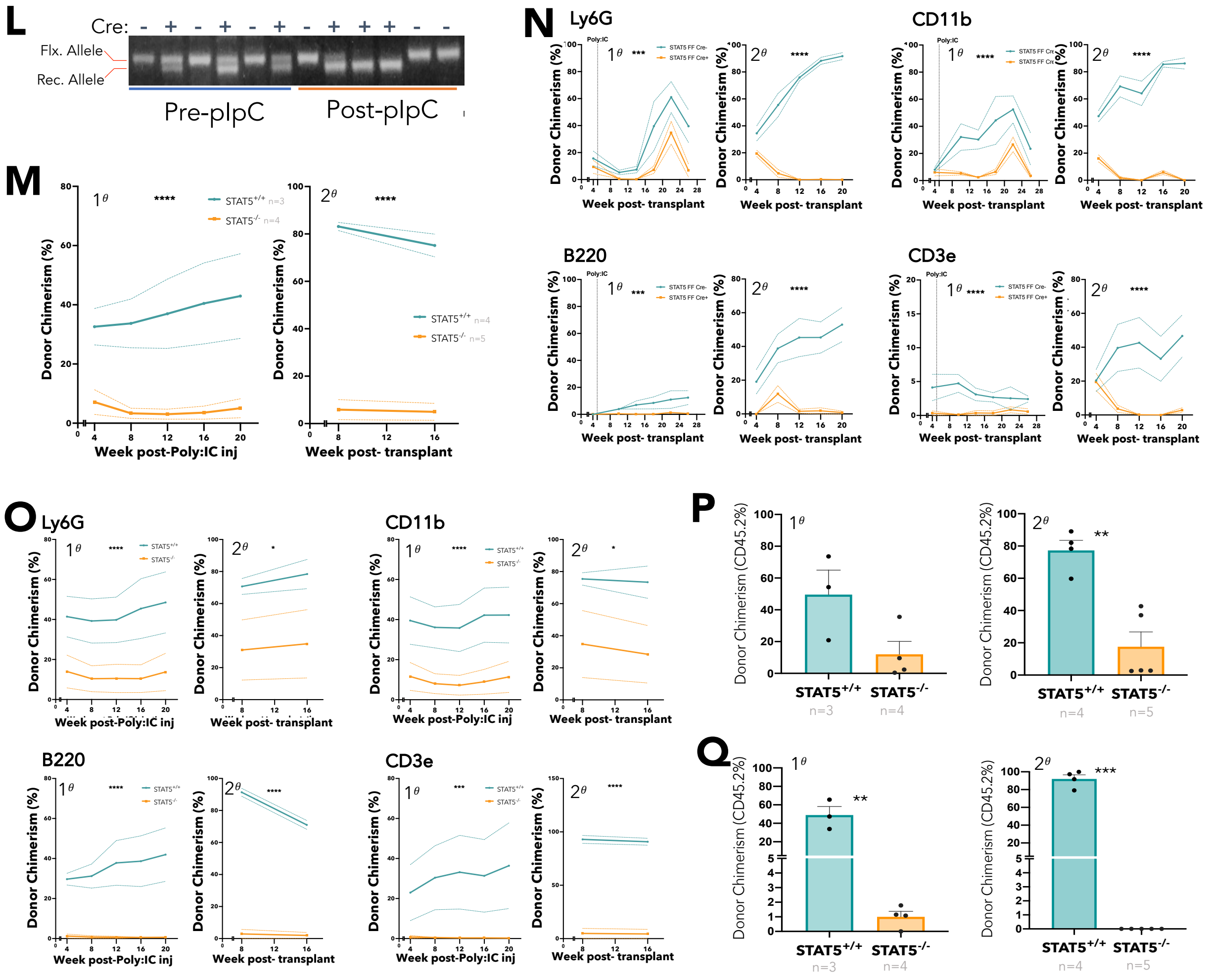


**Supplemental Figure 1: STAT5 loss results in defective HSC function**

(A) Schematic diagram showing induction of STAT5 deletion by repeated Poly:IC treatments. Both STAT5$^{f/f}$Cre$^{+}$ and STAT5$^{f/f}$Cre$^{-}$ mice were treated with Poly:IC over 21 days to induce deletion of *Stat5* locus in Mx1Cre$^{+}$ mice. Further analysis was conducted at least 4 weeks post final injection. (B) Diagram depicting floxed allele with LoxP sites indicated and the recombined allele after Cre mediated deletion of the *Stat5* locus. (C) PCR gel-electrophoresis picture showing recombined band only present in Cre$^{+}$ mice. Comparison of PCR band intensity for floxed and recombined alleles was used to estimate recombination rate. (D) line graph showing RBC, HCT, WBC and PLT counts in peripheral blood from WT and STAT5-deficient mice. Asterisks indicate significant differences by ANOVA column factor (***, P<0.001; *, P<0.001). E) Bar plot showing the number of BMMNCs per 6 hind-leg bones in WT and STAT5-deficient mice (mean ± SEM). (F) Bar plots showing the frequency of MPP1 (Flk2$^{-}$CD150$^{+}$CD48$^{-}$LSK), MPP2 (Flk2$^{-}$CD150$^{+}$CD48$^{+}$LSK), MPP3 (Flk2$^{-}$CD150$^{-}$CD48$^{+}$LSK) and MPP4 (Flk2$^{+}$CD150$^{-}$CD48$^{+}$LSK) cells in BMMNCs from WT and STAT5-deficient mice (mean ± SEM). (G) Bar plots showing the frequency of megakaryocyte (CD41$^{+}$CD42d$^{+}$) and erythroid precursor cells (I, CD71$^{hi}$Ter119$^{mid}$; II, CD71$^{hi}$Ter119$^{hi}$; III, CD71$^{mid}$Ter119$^{hi}$; IV, CD71$^{low}$Ter119$^{hi}$) in BMMNCs (mean ± SEM). (H) Bar plot showing the frequency of monocytes, dendritic and natural-killer cells (Ly6G$^{+}$), granulocytes and macrophages (CD11b$^{+}$), and T-cells (CD3e$^{+}$) in BMMNCs (mean ± SEM). (I) Bar plots showing the frequency of MKp (Lin$^{-}$Sca1$^{-}$cKit$^{+}$CD150$^{+}$CD41$^{+}$), pre-Meg/E (Lin$^{-}$Sca1$^{-}$cKit$^{+}$CD41$^{-}$CD16/32$^{-}$CD105$^{-}$CD150$^{+}$), pre-GM (Lin$^{-}$Sca1$^{-}$cKit$^{+}$CD41$^{-}$CD16/32$^{-}$CD105$^{-}$CD150$^{-}$), GMP (Lin$^{-}$Sca1$^{-}$cKit$^{+}$CD41$^{-}$CD16/32$^{+}$CD150$^{-}$), and pre-CFUe (Lin$^{-}$Sca1$^{-}$cKit$^{+}$CD41$^{-}$CD16/32$^{-}$CD105$^{+}$CD150$^{+}$) cells in BMMNCs (mean ± SEM). (J) Bar plot showing the frequency of monocytes, dendritic and natural-killer cells granulocytes and macrophages, B cells (B220$^{+}$) and T-cells in spleen mono-nuclear cells (mean ± SEM). Asterisks indicate significant differences by Student's t test (*, P<0.05; **, P<0.01; ***, P<0.001), unless otherwise specified. (K) Left; UMAPs showing the density of cells in WT and STAT5-deficent HSPCs from scRNAseq data of FACS sorted LK (Lin$^{-}$c-Kit$^{+}$) cells from WT (n=3) and STAT5-deficient (n=3) BM. Cells were projected onto a combined landscape of over 40,000 HSPCs from Nestorowa *et al.* (2016) and Dahlin *et al.* (2018), and nearest neighbours' analysis was performed to ascribe cell identity and cell type annotation. Cell-type key is shown on bottom left UMAP. Right; bar graph showing the relative abundance of HSPC cell types. (L) Representative PCR gel showing floxed and recombined bands in peripheral bloods of recipients pre- and post- poly:IC. PCR shows partial recombination of the STAT5 locus occurred in Cre$^{+}$ donors in recipients before Poly:IC injections, but near complete recombination is seen after Poly:IC injections. (M) Connected line graphs showing donor chimerism in peripheral blood mono-nuclear cells at each time point in primary and secondary recipients (mean ± SEM). Asterisks indicate significant differences by ANOVA column factor (****, P<0.0001). Results were from an independent replicate competitive transplant experiment (Experiment II). (N) Connected line graphs showing donor chimerism in Ly6G$^{+}$, CD11b$^{+}$, B220$^{+}$ and CD3e$^{+}$ peripheral blood cells in primary and secondary recipients (mean ± SEM). Asterisks indicate significant differences by ANOVA column factor (****, P<0.0001; ***, P<0.001). Results from experiment I. (O) Connected line graphs showing donor chimerism in Ly6G$^{+}$, CD11b$^{+}$, B220$^{+}$ and CD3e$^{+}$ peripheral blood cells in primary and secondary recipients (mean ± SEM). Asterisks indicate significant differences by ANOVA column factor (****, P<0.0001; *, P<0.05). Results from experiment II. (P) Bar plots showing total BMMNC donor chimerism in primary and secondary recipients (mean ± SEM) from experiment II. (Q) Bar plots showing LT-HSC donor chimerism in primary and secondary recipients (mean ± SEM) from experiment II. Asterisks indicate significant differences by Student’s t test (***, p<0.001; **, p<0.01).

***Supplemental Figure 2:*** **STAT5-deficient HSCs display reduced cell cycle entry, increased differentiation, and reduced retention of lineage-negative progeny.**

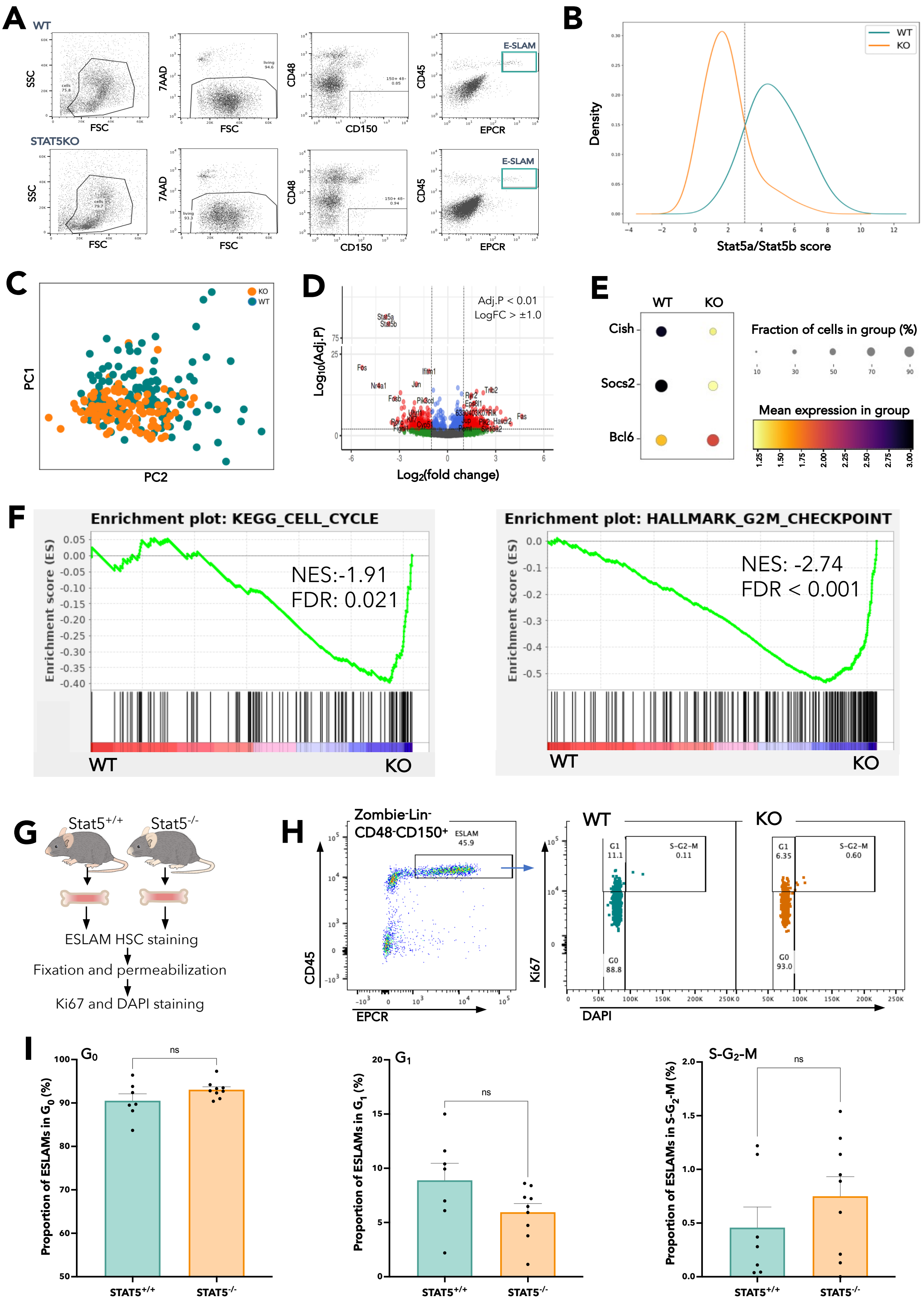

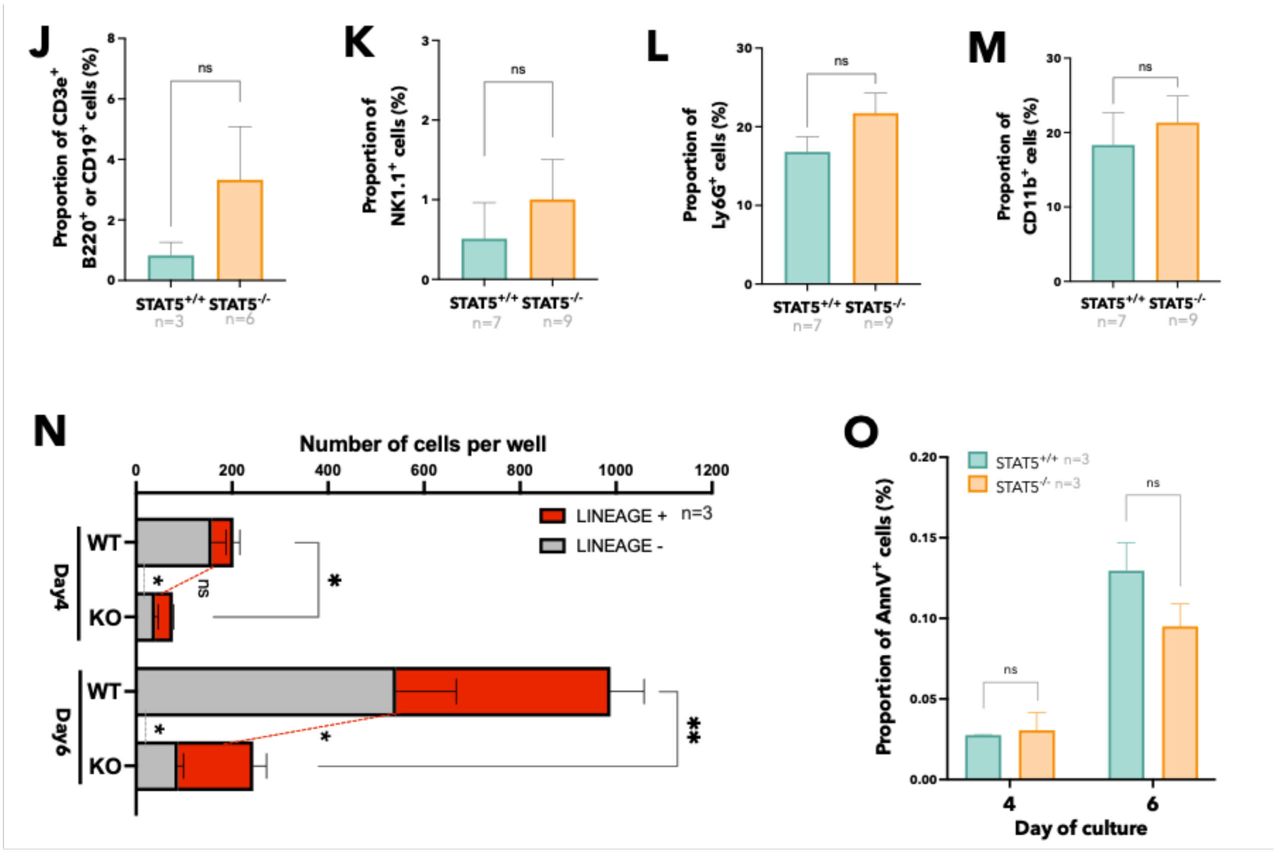


***Supplemental Figure 2:*** ***STAT5-deficient HSCs display reduced cell cycle entry, increased differentiation, and reduced retention of lineage-negative progeny.***

(A) Gating strategy used to isolate ESLAM HSCs from STAT5$^{-/-}$ and WT mice for ScRNAseq. Bone marrow cells were collected from 6 hind-leg bones and lineage depleted before antibody staining for ESLAM markers (CD45$^{+}$CD48$^{-}$CD150$^{+}$ECPR$^{+}$) and FACS sorting. (B) Line graph showing the kernel density estimates of HSCs from STAT5 deficient mice and WT mice showing the geometric mean score of *Stat5a* and *Stat5b* expression. The dotted vertical line indicates where HSCs from WT and STAT5 deficient mice overlap in Stat5a/b transcripts. All HSCs from STAT5 deficient mice above the dotted line were removed before further analysis (n=23 HSCs removed). (C) PCA analysis showing the location of STAT5$^{-/-}$ and WT HSCs based on principal component analysis. (D) Volcano plot showing differentially expressed genes (red dots) in $STAT5^{-/-}$ HSCs with a $Log_2$ Fold Change >1.0 and Benjamini-Hochberg corrected p-value <0.01. (E) Dot plot showing differentially expressed canonical STAT5 target genes. (F) Gene set enrichment analysis (GSEA) plots showing depleted cell cycle related signatures in STAT5-deficient ESLAM HSCs. Normalised enrichment scores (NES) and false discovery rate (FDR) are indicated. (G) Schematic diagram showing cell cycle analysis of HSCs using intracellular flow cytometry after staining with Ki-67 and DAPI. BM from $STAT5^{-/-}$ and WT mice was lineage depleted, stained with ESLAM HSCs panel. Cells were then fixed and permeabilised and stained for Ki67 and DAPI for cell cycle analysis. (H) Example of gating of fixed and permeabilised BM lineage negative cells for ESLAM HSC cell cycle analysis with different stages of cell cycle indicated. (I) Bar plot showing the proportion of ESLAM HSCs in $G_0$, $G_1$ and S-$G_2$-M phases of the cell cycle based on Ki-67 and DAPI staining. Each dot represents the mean of ESLAMs in an indicated phase from one WT or STAT5-deficient mouse. (J) Bar plot showing the frequency of lymphocyte lineage marker ($CD3e^+$/$CD19^+$/$B220^+$) expressing cells at day 5 in *ex vivo* IL-11/SCF culture (mean ± SEM). (K) Bar plot showing the frequency of natural killer lineage marker ($NK1.1^+$) expressing cells (mean ± SEM) produced by WT or STAT5-defcient HSCs at day 5 in *ex vivo* IL-11/SCF culture. (L) Bar plot showing the frequency of monocytes and neutrophils lineage marker ($Ly6G^+$) expressing cells (mean ± SEM) produced by WT or STAT5-defcient HSCs at day 5 in *ex vivo* IL-11/SCF culture. (M) Bar plot showing the frequency of granulocytes and macrophages lineage marker ($CD11b^+$) expressing cells (mean ± SEM) produced by WT or STAT5-defcient HSCs at day 5 in *ex vivo* IL-11/SCF culture. (N) Bar plots showing the number of cells per well at day 4 and 6 produced by 50 WT or STAT5-deficient

ESLAM HSCs in *ex vivo* IL-11/SCF culture. The number of cells expressing lineage markers ($Ter119^+$ $Ly6g^+$ $CD11b^+$ $B220^+$ or $CD3e^+$) are in red; the number of lineage negative cells are in grey (mean ± SEM). (O) Bar plot showing the proportion of AnnexinV+ cells in cultures at days 4 and 6 in *ex vivo* IL-11/SCF culture (mean ± SEM). Asterisks indicate significant differences by Student's t test (*, $p<0.05$; **, $p<0.01$).

***Supplemental Figure 3:*** **Unphosphorylated STAT5 constrains HSC differentiation and upregulates transcriptional programs associated with HSC maintenance.**

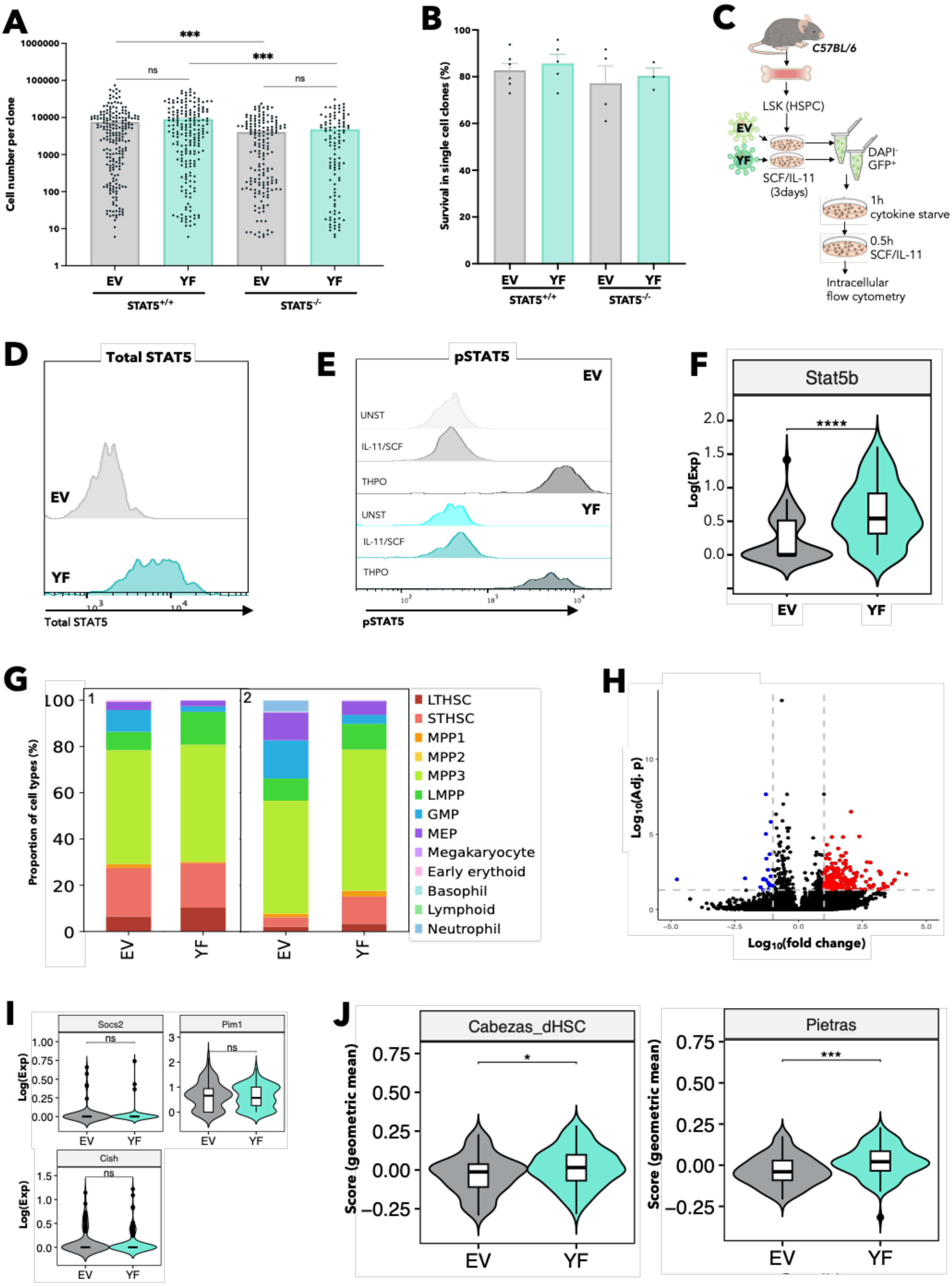

***Supplemental Figure 3:*** **Unphosphorylated STAT5 constrains HSC differentiation and upregulates transcriptional programs associated with HSC maintenance.**

(A) Bar plot showing the number of cells per well at day 7 after single cells were sorted per well into SCF/IL-11 maintenance conditions.[2] Each dot represents a single clone and bars represent mean cell number (± SEM); results were from 4 independent experiments. (B) Bar plots showing clone survival rate after 5 days in culture. Clone survival rate was shown as the proportion of wells that contained cells at day 5. Each dot represents the frequency of surviving clones from each biological replicate; bars show the mean ± SEM survival from all experiments. (C) Schematic diagram showing intracellular flow cytometry analysis of pSTAT5 and total STAT5 protein levels. Lineage$^{-}$Sca1$^{+}$cKit$^{+}$ (LSK) were FACS sorted from WT bone marrow and transduced with lentivirus containing STAT5B-Y699F(YF), or EV in SCF/IL-11 maintenance cultures.[2] After three days, GFP$^{+}$ living cells were sorted and starved in cytokine free media for 1h, followed by a 30-minute stimulation with IL-11/SCF maintenance cultures. After stimulation cells were fixed and permeabilized and stained with anti-STAT5 antibody or anti-pSTAT5 antibody. (D) Example of histograms for total-STAT5 antibody staining of LSK cells transduced with STAT5B-YF or EV lentivirus. GFP+ living cells were sorted 3-days post transduction and starved in cytokine free media for 1h before being placed into either starved media or, IL-11/SCF culture, a media with cytokine cocktail containing high doses of THPO (100ng/ml) was included for positive control. (E) Example of histograms for pSTAT5 antibody staining of LSK cells transduced with STAT5B-YF or EV lentivirus. GFP+ living cells were sorted 3-days post transduction and starved in cytokine free media for 1h before being placed into either starved media or, IL-11/SCF culture, or media containing a cytokine rich cocktail containing high doses of THPO (100ng/ml) was included for positive control. (F) Violin plot showing the normalised expression of *Stat5b*. Asterisks indicate significant differences by Benjamini-Hochberg corrected p-value (****, adj. p<0.001). (G) Bar plots showing the frequency of annotated cell types in each independent experiment after datasets were projected onto a combined landscape of over 40,000 HSPCs from Nestorowa *et al.* (2016) and Dahlin *et al.* (2018). After projection, nearest neighbours analysis was performed to annotate cell types within datasets. (H) Volcano plot showing differentially expressed genes (red dots) in STAT5B-YF HSCs compared to EV HSCs. Cut-off; >1.0 $Log_2$ Fold Change and Benjamini-Hochberg corrected p-value <0.05. (I) Violin plots showing the expression level and frequency of cells expressing canonical pSTAT5 target genes; Cish, Pim1 and Socs2 in STAT5B-YF or EV infected LT-HSCs. (J) Violin plots showing the geometric mean score of STAT5B-YF or EV infected LT-HSCs for two published HSC signatures: dHSC; dormant HSCs compared to active HSCs using retinoic acid reporter mice, in Cabezas-Wallscheid *et al.* (2017)[16] and $Flt3^-$ SLAM; LT-HSCs compared to multipotent progenitors from Pietras *et al.* (2014).[17] Asterisks indicate significant differences by Student's t test (*, p<0.05; ***, p<0.001; ****, p<0.0001).

***Supplemental Figure 4:*** **Unphosphorylated STAT5 enhances HSPC clonogenicity *in vitro* and HSC maintenance *in vivo*.**

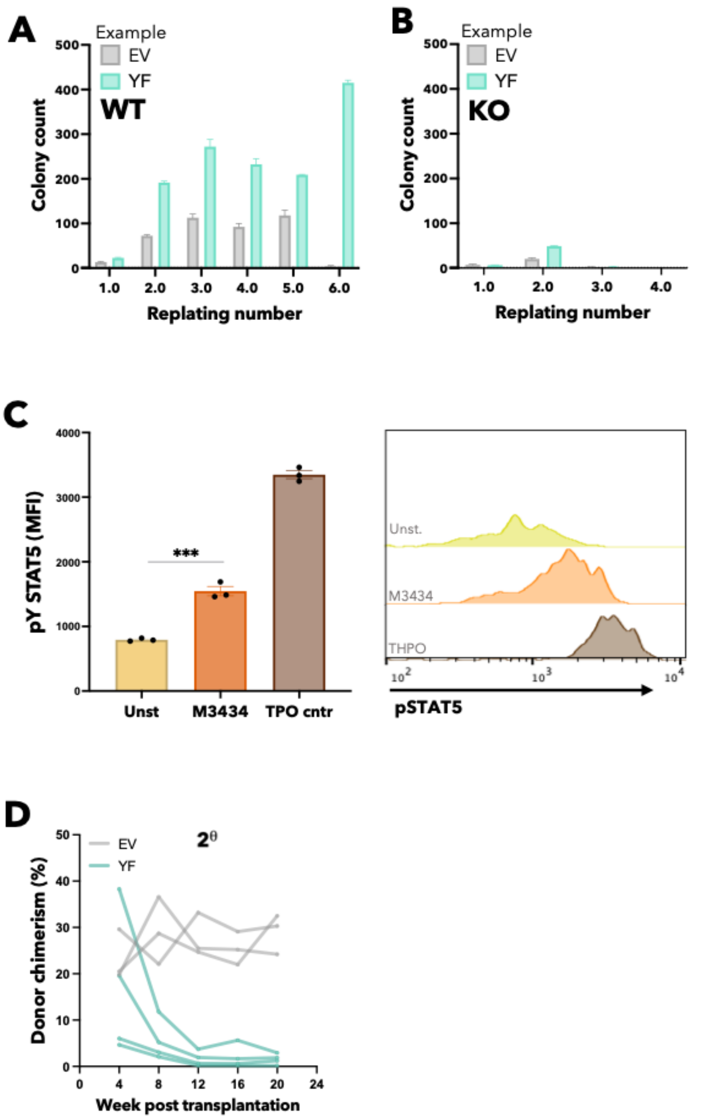


***Supplemental Figure 4:*** **Unphosphorylated STAT5 enhances HSPC clonogenicity *in vitro* and HSC maintenance *in vivo*.**

(A) Bar graph showing an example of the number of colonies at each replating week for one biological replicate of WT HSCs transduced with STAT5B-YF, or EV (n=2 technical replicates). (B) Bar graph showing number of colonies at each replating week for STAT5-deficient HSCs transduced with STAT5B-YF, or EV (n=2 technical replicates). Data was collected at the same time as A. (C) Left; Bar plots showing the mean fluorescent intensity of pSTAT5 antibody staining by intracellular flow-cytometry analysis in ESLAM HSCs unstimulated, semi-solid methylcellulose condition used in replating assays (M3434), and THPO (200ng/ml) positive control conditions (mean ± SEM). Right; example of histograms showing mean fluorescent intensity of pSTAT5 staining of ESLAM HSCs stimulated with indicated culture conditions. Each bar plot shows the mean ±SEM. (D) Line graph showing the peripheral blood donor chimerism of HSCs transduced with EV (n=3), or STAT5B-Y699F (n=4) for each recipient mouse in secondary competitive transplants. Asterisks indicate significant differences by Student's t test (***, $p<0.001$).

***Supplemental Figure 5*** **Ruxolitinib enhances HSPC clonogenicity and maintains transplantable HSCs.**

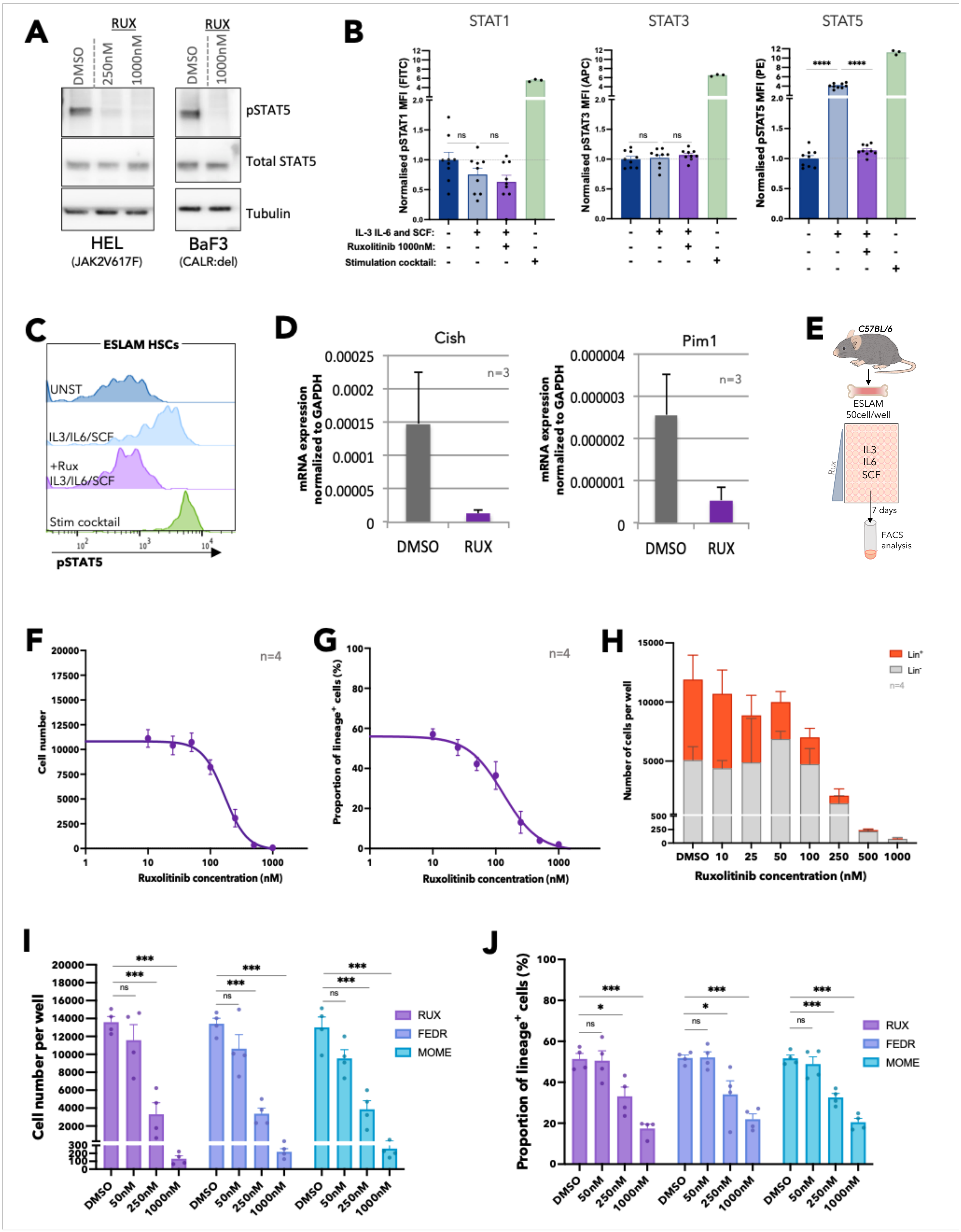

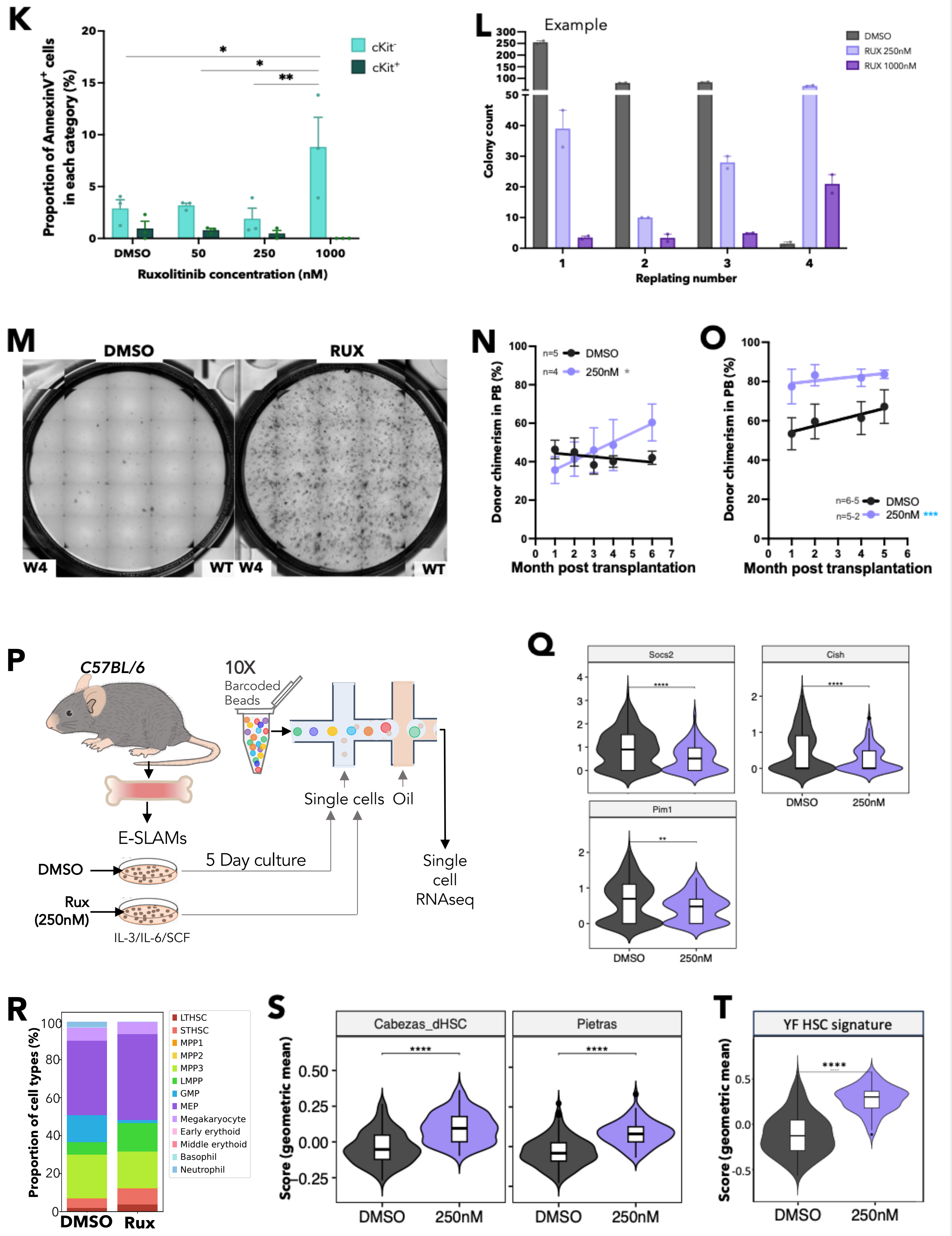
K
Proportion of AnnexinV+ cells in each category (%)
cKit-
cKit+
Ruxolitinib concentration (nM)
DMSO
50
250
1000
L
Example
DMSO
RUX 250nM
RUX 1000nM
Colony count
Replating number
M
DMSO
RUX
W4
WT
N
n=5 DMSO
n=4 250nM
Donor chimerism in PB (%)
Month post transplantation
O
n=6-5 DMSO
n=5-2 250nM
P
C57BL/6
10X
Barcoded
Beads
Single cells
Oil
E-SLAMs
DMSO
Rux (250nM)
5 Day culture
IL-3/IL-6/SCF
Single cell RNAseq
Q
Socs2
Cish
Pim1
DMSO
250nM
R
Proportion of cell types (%)
LTHSC
STHSC
MPP1
MPP2
MPP3
LMPP
GMP
MEP
Megakaryocyte
Early erythoid
Middle erythoid
Basophil
Neutrophil
DMSO
Rux
S
Cabezas_dHSC
Pietras
Score (geometric mean)
T
YF HSC signature

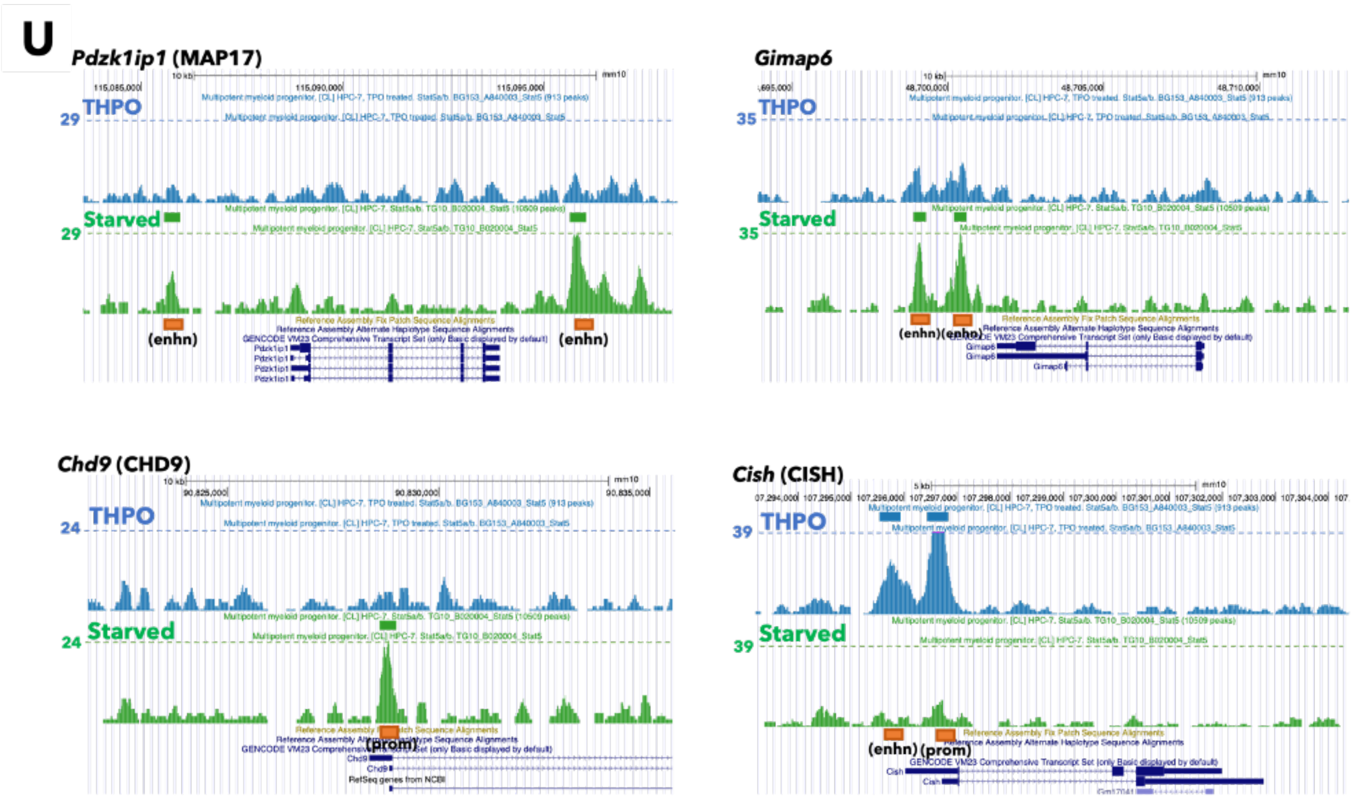


***Supplemental Figure 5:*** **Ruxolitinib enhances HSPC clonogenicity and maintains transplantable HSCs.**

(A) Western blot showing the staining of anti-pSTAT5A/B and anti-total STAT5 antibodies with and without ruxolitinib in HEL cells and Ba/F3 cells expressing both MPL and mutant CALR (52bp deletion, del). (B) Bar plots showing the mean fluorescent intensity of staining of anti-pSTAT1, anti-pSTAT3, and anti-pSTAT5 antibodies by intracellular-flow cytometry in ESLAM HSCs cultured in cytokine free conditions or in cultures stimulated with IL-3/IL-6/SCF[3] cultures ± ruxolitinib, or a media with stimulation cocktail (THPO, Flt3-L and IFN-α). Fluorescent pSTAT staining was normalized to the unstimulated condition for each graph (mean ± SEM). Each dot represents the mean fluorescent intensity of ESLAMs from a single mouse. (C) Example of histograms of pSTAT5 fluorescent staining of ESLAM HSCs cultured in the given culture settings. (D) Bar plots showing the normalised mRNA expression of canonical STAT5 target genes in ESLAM HSCs after 2hrs of exposure to vehicle or ruxolitinib in the presence of SCF/IL-3/IL-6[3] (mean ±SEM). (E) Schematic diagram showing *in vitro* assays of WT HSCs treated with Rux. 50 WT ESLAM HSCs were sorted per well into complete SCF/IL-3/IL-6 media[3] with scaled doses of ruxolitinib or DMSO vehicle and were analysed by flow-cytometry 7 days later. (F) Inhibitor response curves showing the number of cells per well at each dose of ruxolitinib after 7 days in culture (mean ± SEM). Experiment described in ‘E’. (G) Inhibitor-response curves showing the proportion of cells per well that express mature lineage markers (Ter119$^+$/Ly6g$^+$/CD11b$^+$/B220$^+$/CD3e$^+$) after 7 days in culture at different ruxolitinib concentrations (mean ± SEM). Experiment described in ‘E’. (H) Bar plot showing the number of cells that express lineage markers (Lin$^+$) and number of lineage-marker negative (Lin$^-$) cells per well at day 7 (mean ± SEM). (I) Bar plots showing the cell number per well of HSC derived cultures after 7 days in IL-3/IL-6/SCF cultures in the presence of varying doses of ruxolitinib, Fedratinib or Momelotinib, starting from 50 ESLAM HSCs/well (mean ± SEM). (J) Bar plots showing the proportion (%) of lineage positive cells (Ter119$^+$/Ly6g$^+$/CD11b$^+$/B220$^+$/ CD3e$^+$) per well of HSC derived cultures after 7 days in IL-3/IL-6/SCF cultures in the presence of varying doses of ruxolitinib, Fedratinib or Momelotinib, starting from 50 ESLAM HSCs/well (mean ± SEM). (K) Bar plot showing the proportion of apoptotic cKit$^+$ or cKit$^-$ cells (Annexin V$^+$) in cultures derived from lineage depleted BM after 16h with ruxolitinib or vehicle (mean ± SEM). (L) Bar graph showing an example of colony numbers recorded at each replating week for WT HSCs from 1 mouse, treated for 7-days with vehicle or ruxolitinib at 250nM or 1000nM (n=2 technical replicates). (M) Example images of colonies at the final week (4$^{th}$ week) of serial colony plating for WT HSCs treated for 7-days with vehicle or ruxolitinib at 250nM. (N) Scatter dot plot with linear regression line of best fit showing the peripheral blood donor chimerism in primary competitive transplants of 5-day *ex vivo* cultured HSCs with ruxolitinib or vehicle. 50 ESLAMs from WT mice were seeded per well in IL-3/IL-6/SCF culture conditions and given DMSO or 250nM of ruxolitinib for 5 days before the wells were harvested and pooled for each condition and an equivalent of 50 starting ESLAMs was transplanted per recipient with $3x10^5$ competitor bone marrow cells. Each dot indicates mean donor chimerism (mean ± SEM). Grey asterisks indicate significant differences in the slopes of linear regressions modelling ruxolitinib treated donor cell chimerism compared to DMSO treated donor cell chimerism in transplant (*, $p<0.05$). (O) Scatter dot plot with linear regression line of best fit showing the peripheral blood donor chimerism in secondary transplants of 5-day *ex vivo* cultured HSCs with ruxolitinib or vehicle. Each dot indicates mean donor chimerism (mean ± SEM). Blue asterisks indicate significant differences in y-intercepts of linear regressions modelling ruxolitinib treated donor cell chimerism compared to DMSO treated donor cell chimerism in transplants (***, $p<0.001$). (P) Schematic diagram showing experimental outline for scRNAseq of WT ESLAM HSCs treated with RUX or DMSO. WT ESLAM HSCs were FACS sorted and cultured in IL-3/IL-6/SCF complete media[3] with vehicle or 250nM of ruxolitinib and were allowed to expand for 5 days. Living cells were then sorted for 10X Genomics ScRNAseq. (Q) Violin plots showing the mean expression and frequency of LT-HSCs expressing canonical pSTAT5 target genes; Cish, Pim1, and Socs2. LTHSCs were transcriptionally defined within scRNAseq dataset of the day-5 HSC derived cultures treated with DMSO or ruxolitinib. (R) Bar plots showing the frequency of annotated cell types after datasets were projected onto a combined landscape from Nestorowa *et al.* (2015) and Dahlin *et al.* (2018). Nearest neighbours analysis was performed after projection to assign cell types. (S) Violin plots showing the distribution of geometric mean scores (mean of transcript counts across genes within a given dataset) of DMSO or Ruxolitinib treated LT-HSCs for two previously published HSC signatures: dHSC (dormant HSCs) compared to active HSCs from Cabezas-Wallscheid *et al.* (2017)[16] and Flt3$^-$SLAM LT-HSCs compared to multipotent progenitors from Pietras *et al.* (2014).[17] (T) Violin plots showing the geometric mean scores (mean of transcript counts across genes within dataset) of DMSO or Ruxolitinib treated LT-HSCs against the signature of STAT5-YF LT-HSCs from Figure 3. This signature was derived by taking the top 100 significantly upregulated genes (FDR <0.01) in STAT5-YF expressing LT-HSCs compared to control EV transduced LT-HSCs from Figure 3. (U) Representative histogram plots showing the presence of STAT5 binding near example genes known to promote HSC maintenance; these include Pdzk1ip1 (MAP17), Chd9, and Gimap6 in HPC7 cells in cytokine starved condition (i.e. uSTAT5 peak in green), whereas no STAT5 binding is detected in cells cultured with 100ng/ml of THPO (pSTAT5 peak in blue).[18] Cish is provided as an example of a pSTAT5 target gene, i.e. STAT5 binding is detected in THPO stimulated conditions, and is absent in cytokine starved conditions. Asterisks indicate significant differences by Student's t test (****, p<0.0001; ***, p<0.001; **, p<0.01; *, p<0.05) unless otherwise indicated.

***Supplementary Figure 6: Ruxolitinib maintains murine and human myeloproliferative neoplasm HSPCs.***

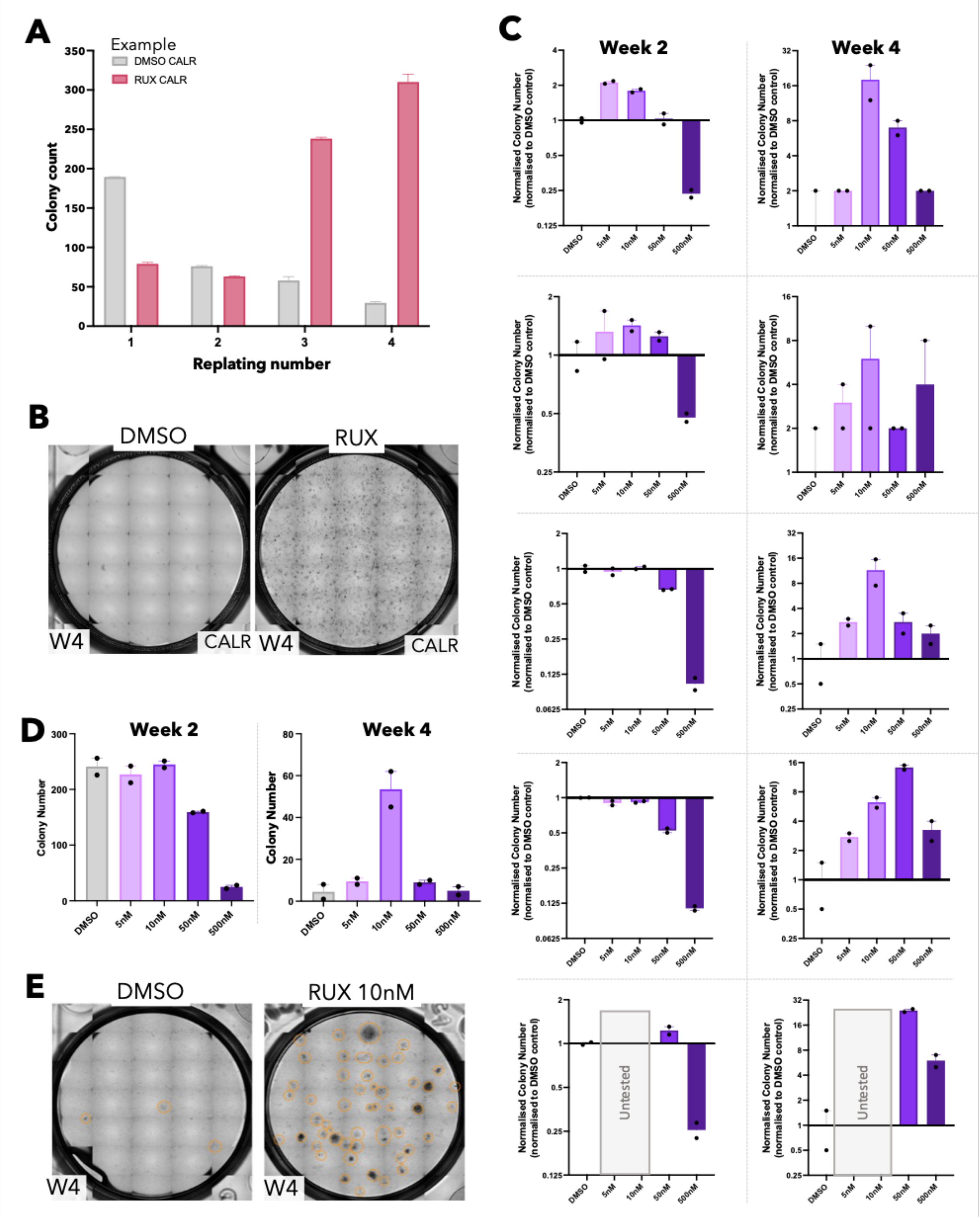

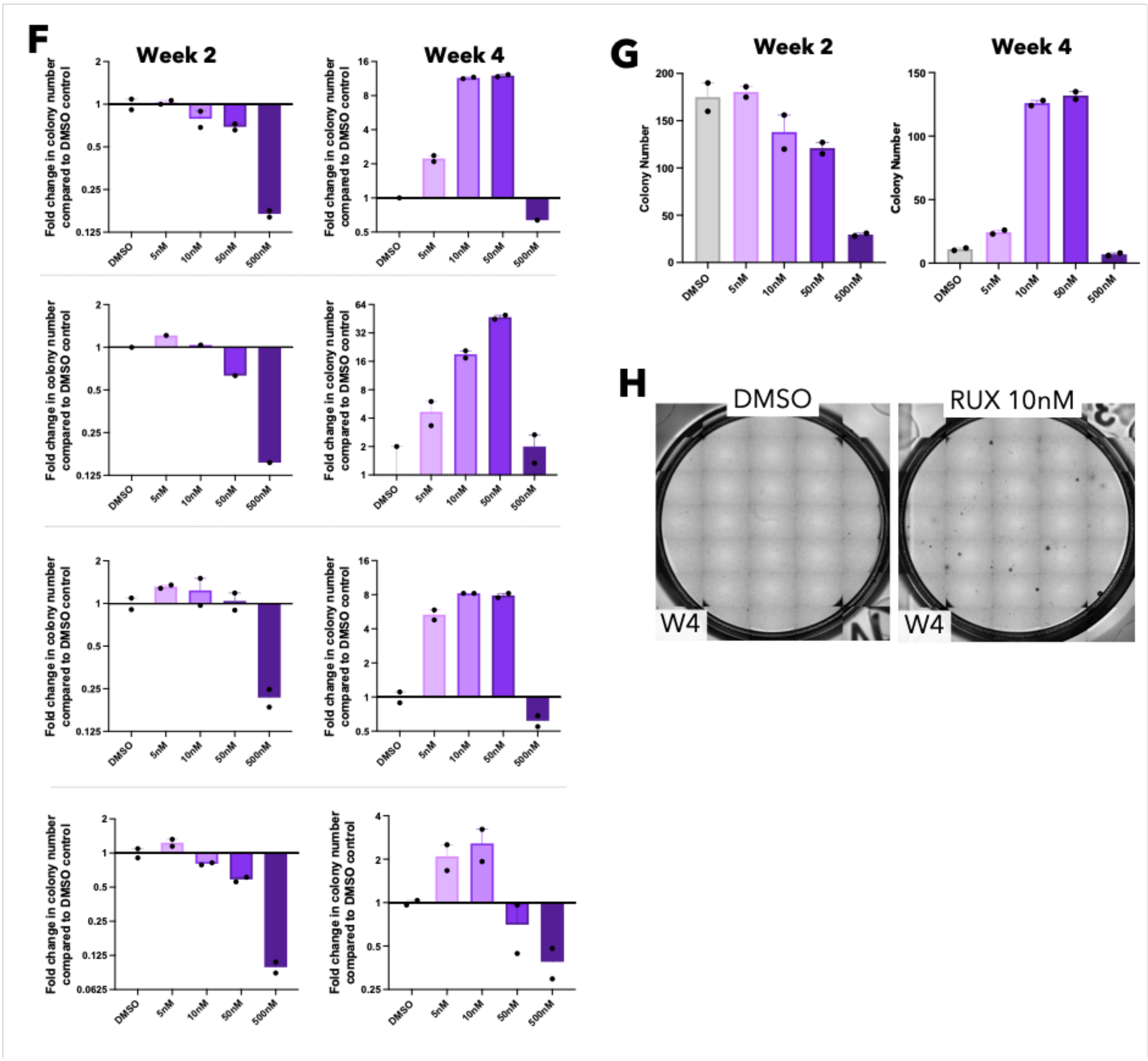


***Supplementary Figure 6: Ruxolitinib maintains murine and human myeloproliferative neoplasm HSPCs.***

(A) Bar graph showing colony numbers recorded at each replating week for CALR homozygous mutant HSCs from CALRdel/del mice. ESLAM HSCs were treated for 7-days with vehicle or ruxolitinib at 250nM (n=2 technical replicates). (B) Example of colony images showing the number and size difference of the colonies at the final (fourth) week of replating by CALR homozygous mutant HSCs treated with DMSO or ruxolitinib for 7 days in IL-3/IL-6/SCF cultures. (C) Bar plots showing the number of colonies produced by human HSCs ($CD34^{+}CD38^{-}CD45RA^{-}$) cultured for 7 days in the presence of ruxolitinib, normalised to the number of colonies produced by HSCs cultured for 7 days with DMSO (fold change). Each row of the bar graphs represents data from an individual donor. Left bar graph showing the fold-change in colony numbers in the first round of colony formation (2 weeks in methylcellulose). Right bar graph showing fold-change colony numbers in the second round of colony formation (4 weeks in methylcellulose). HSCs from 4 individual healthy donor were treated with DMSO, 5nM, 10nM, 50nM, or 500nM of ruxolitinib, which correspond to Figure 6G in the main text. A $5^{th}$ healthy donor was included, which was from a preliminary test where fewer doses of ruxolitinib were tested (50nM and 500nM; bottom row). (D) Bar plots showing the colony numbers (mean ± SEM; n=2 technical replicates shown as dots) at the first plating (left) and the second plating (right), corresponding to normalised data in *Supl. 6C*. (E) Example images showing number and size of colonies produced by healthy human donor HSCs cultured with DMSO or ruxolitinib for 7 days, at the final plating (4-weeks). (F) Bar plots showing the number of colonies produced by HSCs ($CD34^{+}CD38^{-}CD45RA^{-}$) that were sorted from human myelofibrosis patient and cultured for 7 days in the presence of ruxolitinib, data were normalised to the number of colonies produced by HSCs cultured for 7 days with DMSO. Each row represents data from an individual donor. Left graph showing normalised colony numbers (n=2 technical replicates) in the first round of colony formation (2 weeks in methylcellulose). Right graph showing normalised colony numbers in the second round of colony formation (4 weeks in methylcellulose). (G) Bar plots showing the colony numbers (mean ± SEM; n=2 technical replicates) at the first plating (left) and the second plating (right), corresponding to normalised data in F. (H) Example images showing size and number of colonies produced by myelofibrosis patient HSCs cultured with DMSO or ruxolitinib for 7 days, at the final plating (4-weeks).

#### Table legends

**Table 1: Differentially expressed genes in STAT5-deficient HSCs.** ESLAM HSCs from WT and STAT5-deficient ESLAM mice were FACS isolated from hind leg bones before plate based scRNA sequencing by Smartseq2 platforms. The transcriptomes of WT and $STAT5^{-/-}$ ESLAMs were compared by differential gene expression (DEx) analysis. All detected differentially expressed genes are listed.

**Table 2: Significantly enriched and depleted pathways in STAT5 deficient HSCs.** Significantly enriched or depleted gene sets in STAT5-deficient ESLAM HSCs from Hallmarks and KEGG databases as determined by gene set enrichment analysis (GSEA) from ranked differentially expressed gene lists. Gene sets with an FDR q-value of less than 0.25 were considered significantly enriched in STAT5 deficient HSCs.

**Table 3: Differentially expressed genes in uSTAT5B overexpressing transcriptionally defined LT-HSCs.** WT ELSAM HSCs were transduced with virus and cultured for 5 days in IL-11/SCF conditions[2] before 10X scRNA sequencing. The transcriptomes of transcriptionally isolated LT-HSCs infected with lentivirus containing STAT5B-YF were compared to the transcriptomes of EV transduced LT-HSCs. Differential gene expression (DEx) analysis was performed. All detected differentially expressed genes are listed.

**Table 4: Differentially expressed genes in ruxolitinib treated transcriptionally defined LT-HSCs.** WT ESLAM HSCs were cultured in IL3/IL-6/SCF conditions for 5 days with DMSO or ruxolitinib (250nM) before 10X scRNA sequencing. The transcriptomes of transcriptionally isolated LT-HSCs treated with ruxolitinib were compared to the transcriptomes of LT-HSCs treated with DMSO. Differential gene expression (DEx) analysis was performed. All detected differentially expressed genes are listed.

**Table 5: Colony counts from serial replating of human healthy donor HSCs cultured with ruxolitinib or DMSO.** Healthy human platelet apheresis donor HSCs were harvest from apheresis filters and placed into high cytokine cultures (EXPER[15]) for 7 days with ruxolitinib (5nM, 10nM, 50nM or 500nM) at 400 cells per well. Cells were harvested and placed into serial colony replating assays. After 2 weeks, colonies were counted and well contents were harvested and diluted for further colony formation; colonies were counted after 2 weeks.

**Table 6: Colony counts from serial replating of myelofibrosis donor HSCs cultured with ruxolitinib or DMSO.** Circulating HSCs were harvest from the peripheral blood of myelopoiesis patients and placed into high cytokine cultures (EXPER[15]) for 7 days with ruxolitinib (5nM, 10nM, 50nM or 500nM) at 400 cells per well. Cells were harvested and placed into serial colony replating assays. After 2 weeks, colonies were counted and well contents were harvested and diluted for further colony formation; colonies were counted after 2 weeks.

#### References


1. Balazs AB, Fabian AJ, Esmon CT, Mulligan RC. Endothelial protein C receptor (CD201) explicitly identifies hematopoietic stem cells in murine bone marrow. *Blood*. 2006;107(6):2317–2321.
2. Kent DG, Dykstra BJ, Cheyne J, Ma E, Eaves CJ. Steel factor coordinately regulates the molecular signature and biologic function of hematopoietic stem cells. *Blood*. 2008;112(3):560–567.
3. Daniel P, Jung PH, Sam W, et al. The stem/progenitor landscape is reshaped in a mouse model of essential thrombocythemia and causes excess megakaryocyte production. *Sci Adv*. 2022;6(48):eabd3139.
4. Zerbino DR, Achuthan P, Akanni W, et al. Ensembl 2018. *Nucleic Acids Res*. 2018;46(D1):D754–D761.
5. Wu TD, Nacu S. Fast and SNP-tolerant detection of complex variants and splicing in short reads. *Bioinformatics*. 2010;26(7):873–881.
6. Anders S, Pyl PT, Huber W. HTSeq—a Python framework to work with high-throughput sequencing data. *bioinformatics*. 2015;31(2):166–169.
7. Brennecke P, Anders S, Kim JK, et al. Accounting for technical noise in single-cell RNA-seq experiments. *Nat Methods*. 2013;10(11):1093–1095.
8. Ritchie ME, Phipson B, Wu DI, et al. limma powers differential expression analyses for RNA-sequencing and microarray studies. *Nucleic Acids Res*. 2015;43(7):e47–e47.
9. Wolf FA, Angerer P, Theis FJ. SCANPY: large-scale single-cell gene expression data analysis. *Genome Biol*. 2018;19(1):1–5.
10. Love MI, Huber W, Anders S. Moderated estimation of fold change and dispersion for RNA-seq data with DESeq2. *Genome Biol*. 2014;15(12):1–21.
11. Cumaraswamy AA, Lewis AM, Geletu M, et al. Nanomolar-Potency Small Molecule Inhibitor of STAT5 Protein. *ACS Med Chem Lett*. 2014;5(11):1202–1206.
12. Subramanian A, Tamayo P, Mootha VK, et al. Gene set enrichment analysis: a knowledge-based approach for interpreting genome-wide expression profiles. *Proceedings of the National Academy of Sciences*. 2005;102(43):15545–15550.
13. Dahlin JS, Hamey FK, Pijuan-Sala B, et al. A single cell hematopoietic landscape resolves eight lineage trajectories and defects in Kit mutant mice. *Blood*. 2018;blood-2017-12-821413.
14. Nestorowa S, Hamey FK, Pijuan Sala B, et al. A single-cell resolution map of mouse hematopoietic stem and progenitor cell differentiation. *Blood*. 2016;128(8):e20–e31.
15. Belluschi S, Calderbank EF, Ciaurro V, et al. Myelo-lymphoid lineage restriction occurs in the human haematopoietic stem cell compartment before lymphoid-primed multipotent progenitors. *Nat Commun*. 2018;9(1):4100.
16. Cabezas-Wallscheid N, Buettner F, Sommerkamp P, et al. Vitamin A-Retinoic Acid Signaling Regulates Hematopoietic Stem Cell Dormancy. *Cell*. 2017;169(5):807-823.e19.
17. Pietras EM, Lakshminarasimhan R, Techner J-M, et al. Re-entry into quiescence protects hematopoietic stem cells from the killing effect of chronic exposure to type I interferons. *Journal of Experimental Medicine*. 2014;211(2):245–262.
18. Park HJ, Li J, Hannah R, et al. Cytokine-induced megakaryocytic differentiation is regulated by genome-wide loss of a uSTAT transcriptional program. *EMBO J*. 2016;
